# Molecular basis of delayed leaf senescence induced by short-term treatment with low phosphate in rice

**DOI:** 10.64898/2026.01.23.701354

**Authors:** Héctor Martín-Cardoso, Mireia Bundó, Antoni Garcia-Molina, Blanca San Segundo

## Abstract

Leaf senescence is a programmed plant developmental process that can also be regulated by environmental factors, like nutrient availability. Although phosphorus is an essential element determining plants’ growth and productivity, mechanisms underlying adaptation to phosphorus availability in plants are not well understood. In this study, we combined physiological, biochemical and molecular approaches to investigate the effect of phosphate supply on leaf senescence in rice. We show that short-term treatment of rice seedlings with low phosphate increases photosynthetic pigments content, confers tolerance to methyl viologen-induced oxidative stress in chloroplasts, and increases antioxidant enzyme activities. Leaves from low-Pi-treated plants also showed a reduction in membrane lipid peroxidation and electrolyte leakage. Opposite trends were observed in seedlings under high Pi supply, in which accelerated leaf senescence occurs. Further analyses indicated that CRISPR/Cas9-mediated editing of *MIR827*, and subsequent reduction in Pi content, promotes delayed leaf senescence, while Pi accumulation in *MIR827* or *MIR399* overexpressing plants accelerates senescence. These findings strongly support that short treatment with low phosphate delays rice leaf senescence. Transcriptomic analysis demonstrated multiple biological processes underlying adaptation of rice plants to low phosphate, including senescence-associated and metabolic processes. These findings provide novel insights into leaf senescence potentially contributing to sustainable rice production.

## 1. Introduction

Leaf senescence in adult plants is a genetically programmed phenomenon primarily associated with the process of aging in plants. During the senescence period, leaves undergo coordinated changes at the physiological, biochemical and molecular levels aiming to ensure plant survival and reproductive success (Mayta et al., 2019). Leaf senescence can also be induced by environmental stresses such as drought, salinity, high light, or high temperature (Tan et al., 2023). The timing of leaf senescence must be strictly controlled to avoid premature senescence while allowing activation only under unfavourable conditions. In agriculture, delayed senescence (stay green) allows the plant to prolong its photosynthetic capacity and carbon fixation to increase crop yield (Lee and Masclaux-Daubresse, 2021). A fine interplay between developmentally programmed senescence and responses to environmental factors is then crucial for maximizing crop yield and improve survival chances.

One of the most important events in senescent leaves is yellowing due to chlorophyll breakdown. Along with this, delayed senescence extends the functional life of leaves leading to increased photosynthesis, crop yield and stress tolerance. Conversely, premature leaf senescence involves rapid chlorophyll breakdown that negatively impacts crop yield by reducing photosynthesis. Increased ROS production is also observed in leaves during natural senescence and response to environmental stresses (Moustakas et al., 2023). Being chloroplasts one of the most important producers of ROS in plant cells, preventing ROS overproduction and maintenance of redox balance in chloroplast is crucial for normal leaf senescence. Excessive ROS in chloroplasts can directly damage macromolecules (e.g., lipids, DNA, RNA, and proteins) and cause membrane damage thereby damaging the photosynthetic apparatus (Hasanuzzaman et al., 2020). To avoid oxidative damage caused by elevated ROS levels, plants possess scavenging and antioxidant systems that maintain an appropriate balance of ROS in chloroplasts. Further, carotenoids participate in ROS scavenging in chloroplasts.

At the molecular level, leaf senescence is associated to changes in the expression of genes conceptually known as Senescence-Associated Genes (SAGs) with diverse functions, including photosynthesis, oxidative stress, transcriptional regulation and signal transduction, nutrient remobilization and degradation of macromolecules (Woo et al., 2019; Lee and Masclaux-Daubresse, 2021). Furthermore, SAG genes might function as positive or negative regulators of processes related to leaf senescence. Although much of our knowledge on senescence-associated processes comes from studies in the model plant *Arabidopsis thaliana*, research on leaf senescence in crop species has attracted much attention in recent years because of its potential application to improve crop yield (Woo et al., 2019).

In addition to aging, nutrient availability can also dictate the extend and timing of senescence. It is generally believed that high nutrient availability can delay leaf senescence to maximize growth, while low nutrient availability, especially nitrogen, might trigger early senescence to allow for nutrient recycling and remobilization from older tissues to support new growth, a crucial survival strategy (Zhang et al., 2024b). Then, molecular players that function in plant responses to nutrient availability might also participate in senescence-related pathways, an aspect that hasn’t received much attention.

Phosphorus (P), a vital nutrient for plant growth, is typically taken up by plants as inorganic phosphate (Pi) from the soil. Despite its abundance in the environment, P bioavailability is limited due to organic complexation and precipitation with metal cations, leading to poor solubility (George et al., 2016). To overcome Pi limitation, plants implemented adaptive mechanisms to increase Pi uptake from the soil, such as the activation of the so-called Phosphate Starvation Response (PSR) (Puga et al., 2024; Yang et al., 2024; Chiang et al., 2025). Two distinct microRNAs (miRNAs), miR399 and miR827, are key players in the PSR for the control of phosphate (Pi) homeostasis in plants (Chiou et al., 2006; Lin et al., 2013; Puga et al., 2024; Yang et al., 2024). Upon Pi starvation, *MIR399* and *MIR827* expression is induced for down-regulation of the corresponding target genes and enhanced Pi uptake (Puga et al., 2024; Yang et al., 2024). In rice, *MIR399* overexpression and *MIR827* overexpression results in Pi accumulation, whereas CRISPR/Cas9-mediated mutagenesis of *MIR827* causes a reduction in Pi content (Campos-Soriano et al., 2020; Bundó et al., 2024).

In other studies, nutrient imbalances have emerged as critical stressors reducing photosynthetic efficiency and reproductive success in rice (Lu et al., 2025). Being an essential nutrient, both Pi deficiency and excess, might have a substantial impact on plant life dynamics causing significant effects on rice production. However, the impact of stress resulting from inadequate Pi nutrition on physiological and molecular processes in the rice plant, remains poorly explored. Particularly, limited information is available on how Pi content can affect senescence-associated processes in rice.

In this work, we investigated the effect of Pi treatment on leaf senescence in rice plants early during vegetative growth. Physiological, biochemical, and molecular changes were examined in rice seedlings grown under different Pi regimes, which included photosynthetic pigments content, membrane lipid peroxidation and electrolyte leakage, response to methyl viologen-induced oxidative stress, activity of enzymatic antioxidant systems, and transcriptome analysis. Specifically, short-term treatment with low Pi causes a delay in leaf senescence, whereas treatment with high Pi accelerates leaf senescence in rice seedlings. Silencing *MIR827* expression in CRISPR-Cas9-edited plants, and subsequent reduction in Pi accumulation, delays in leaf senescence in rice. Conversely, over-accumulation of Pi in miR399 and miR827 overexpressor plants promotes leaf senescence. Collectively, our results support that treatment of rice plants with low Pi regulates the course of leaf senescence during early growth in rice plants. Furthermore, the molecular framework underlying delayed leaf senescence in response to short treatment with low Pi is presented. A better knowledge of factors affecting delayed leaf senescence during vegetative growth is essential for developing varieties with improved P nutrition capable of thriving Pi limiting conditions with reduced dependency on Pi fertilisers.

## 2. Materials and Methods

### 2.1. Plant material, growth conditions and measurement of Pi content

Rice plants (*Oryza sativa*, *japonica* cv Nipponbare) were grown at 28°C/25°C with a 14h/10h light/dark cycle. The rice cultivars Nipponbare and Tainung 67 (TN67) were used in this study. Production, characterization and growth conditions of miR827 overexpressor and CRISPR/Cas9 mutant plants for *MIR827* (Nipponbare background), and miR399 overexpressor plants (TN67 background) were previously described (Campos-Soriano et al., 2020; Bundó et al., 2024).

For Pi treatment, plants were grown in 50% turface and vermiculite combined with 50% quartz sand. To obtain a homogeneous growth of all the plants, seeds were pregerminated in water for 7 days, transplanted to soil and then fertilized with Hoagland half-strength solution containing the desired Pi concentration, namely 0.025 mM (Low Pi condition), 1.0 mM (Control condition) and 2.5 mM Pi (High Pi condition). The final Pi concentration was adjusted using KH_2_PO_4_. Plants were grown under controlled conditions for 14, 21 and 28 days.

The Pi content of rice leaves was determined as previously described (Ames, 1966; Versaw and Harrison, 2002). Three independent experiments were carried out. Four biological replicates each one from a pool of four different plants, and three technical replicates for each biological replicate were analysed.

### 2.2. Dark-induced leaf senescence assays

Dark-induced leaf senescence (DILS) was investigated on soil-grown rice seedlings at the whole-plant level. For this, rice seedlings were grown as described above (pre-germination for 1 week followed by Pi treatment for 2 weeks) under photoperiod conditions. Seedlings continued growth for 1 week more either in the dark, or under photoperiod condition (control plants). During this additional week, plants were fertilized with the corresponding Pi concentration.

DILS assays on detached leaves, either entire leaves or leaf segments from detached leaves (2-3 cm in length), were also performed. Leaves at position 1 (youngest fully expanded leaf) and position 2 were examined. Entire leaves or leaf segments (2-3 cm in length) were floated in sterilized distilled water (100 mL and 25 mL, respectively) and placed in the darkness at 28°C. The progression of senescence was followed with time. The percentage of leaf senescent area in entire leaves was quantified using APS Assess 2.0 program. For each leaf, leaf area was considered from 72 to 125 pixel colour (red to green, respectively) and senescent area was quantified from 72 to 101 pixel colour (red to orange/yellow, respectively). Three independent experiments were carried out, each one with 20 biological replicates per treatment/genotype.

### 2.3. Measurement of photosynthetic pigment content

Chlorophyll and carotenoid concentrations were measured by extraction of the leaf pigments as previously described (Barja et al., 2021) followed by high-performance liquid chromatography (HPLC) using an Agilent 12000 series HPLC system (Agilent Technologies). Elution of chlorophylls and carotenoids was monitored using a photodiode array detector. Peak areas of chlorophylls (650 nm) and carotenoids (470 nm for lutein, β-carotene, 9-cis-β-carotene, violaxanthin, neoxanthin and canthaxanthin) were determined using the Agilent ChemStation software. Quantification was performed by comparison with commercial standards (Sigma). Three independent experiments were performed, each one with four biological replicates per condition (consisting of 4 plants each replicate).

Chlorophylls and total carotenoid content was also determined spectrophotometrically (Lichtenthaler and Buschmann, 2001). For this, the leaf material (30 mg) was grinded, and chlorophylls were extracted using ethanol 96% (v/v) for 2h at 50°C. Samples were centrifuged for 1 min at 12,000 *g* to remove the cell debris. The content of photosynthetic pigments was determined by measuring the absorbance of the supernatant at 649 and 664 nm for chlorophylls, and 470 nm for carotenoids (Spectramax M3 reader, Molecular Devices, USA) and expressed in mg/g fresh weight (FW) (Lichtenthaler and Buschmann, 2001). Three independent experiments were carried out, each one with four or five biological replicates per condition (consisting of 3 or 4 plants each replicate) and three technical replicates for each biological replicate.

### 2.4. Electrolyte leakage and lipid peroxidation assays

Electrolyte leakage (EL) analyses were carried out as previously described (Campo et al., 2014). Briefly, leaf pieces (2-3 cm segments) were washed in autoclaved Milli-Q water and incubated for 2h in autoclaved Milli-Q water in a shaker. Next, electroconductivity was measured (EC1). The leaf segments were then autoclaved for 20 min and EC2 was measured. The EL was calculated as (EC1/EC2) x 100. Electroconductivity was measured using a conductivity meter (Mettler-Toledo, FiveGo3 Series, Greiefensee, Switzerland). The level of lipid peroxidation was determined by measuring the amount of MDA produced using the thiobarbituric acid reagent (Martín-Cardoso et al., 2024). Absorbance was measured at 532 nm and 600 nm using Spectramax M3 reader (Molecular Devices, USA). Three independent experiments were carried out, each one with four biological replicates per condition (consisting of 6 plants each replicate), and three technical replicates per biological replicate.

### 2.5. Treatment with methyl viologen

Methyl Viologen assays were carried out as previously described (Salvador-Guirao et al., 2019). Leaf segments (2-3 cm in length) were treated with methyl viologen solution (10 µM) for 12 hours in the dark at room temperature and then maintained at 28°C for 3 days under a 16h/8h photoperiod. Chlorophylls and carotenoids were extracted and quantified spectrophotometrically as described above. Pictures were taken with a Leica DM6 microscope under bright field illumination. Three independent experiments were carried out, each one consisting of three biological replicates per condition (4 plants each replicate) and three technical replicates for each biological replicate.

### 2.6. Enzymatic activities

For the analysis of enzymatic antioxidant activity, crude enzyme solution was extracted as described by Zhang et al. (2016). Briefly, rice leaves (0.3 g) were grounded to powder using liquid nitrogen and then homogenized in 3mL of cold 50mM PBS (pH 7.8) containing 0.2 mM EDTA. Homogenates were then centrifuged at 15,000 *g* at 4°C for 20 min and the supernatant was saved for enzyme activity analysis. Protein concentration was determined using the Bradford assay (Bradford, 1976).

APX activity was determined by measuring the decline in absorbance at 290 nm for 3 min (Nakano and Asada, 1981). The reaction assay mixture (1 mL) contained 50 mM potassium phosphate buffer (pH 7.0), 0.5 mM ascorbate, 0.1 mM H_2_O_2_, 0.1 mM EDTA and 15 μL of plant extract.

CAT activity was determined by measuring the decline in absorbance at 240 nm for 5 min (Poli et al., 2018). The reaction assay mixture (1.5 mL) contained 0.75 mL of 100 mM potassium phosphate buffer (pH 7.0), 0.25 mL of 30 mM H_2_O_2_, 25 μL of plant extract and 0.475 mL of distilled water.

POD activity was determined by measuring the increase in absorbance at 470 nm for 5 min (Poli et al., 2018). The reaction assay mixture (3 mL) contained 1 mL of 60 mM potassium phosphate buffer (pH 6.1), 0.5 mL of 16 mM guaiacol, 0.5 mL of 2 mM H_2_O_2_, 15 μL of plant extract and 0.9 mL of distilled water.

Absorbance of enzymatic activity assays was measured using Spectramax M3 reader (Molecular Devices, USA). Three independent experiments were performed. Five biological replicates each one from a pool of 6 different plants, and three technical replicates for each biological replicate were analysed.

### 2.7. Transcriptome Analysis

Total RNA was extracted from rice leaves of 3-week-old plants that had been treated with different Pi supplies for 14 days using Maxwell RSC Plant RNA Kit (Promega). Three biological replicates for Low and Control Pi were examined, each biological replicate consisting of leaves from five individual plants (the youngest totally expanded leaf). RNA quality and integrity were evaluated using an Agilent 2100 Bioanalyzer (Agilent Technologies, Inc.), and only samples with an RNA integrity number (RIN) ≥ 7 were used. RNA libraries were prepared using Hieff NGS Ultima Dual-mode mRNA Library Prep Kit for Illumina (Yeasen, USA) following manufacturer’s instructions and sequenced using Illumina NovaSeq 6000 (Illumina, USA). RNASeq datasets were trimmed using *fastp* (v0.21) (Chen et al., 2018), allowing minimum read length of 35bp and a minimum sequence quality of 25. Read alignment was performed using *STAR* (v2.7.9) with default parameters and the *Oryza sativa* cv *Nipponbare* reference genome (IRGSP-1.0). Read counts were obtained using *featureCounts* (v1.6.0). Counts were filtered for low-expressed transcripts (< 40 counts in at least 3 samples). DEGs were declared based on absolute log_2_ FC ≥ 0.5 to control Pi (adjusted *P* ≤ 0.01, Wald’s test). GO enrichment analysis was performed using the GO webtool (adjusted *P* ≤ 0.05; Fisher’s Exact test) (https://www.geneontology.org) and non-redundant terms were selected and plotted using ReviGO (https://www.revigo.irb.hr/).

### 2.8. Expression analysis by RT-qPCR

Total RNA was extracted using the Maxwell RSC Plant RNA Kit (Promega). Total RNA (1 μg) was retrotranscribed using the High Capacity cDNA reverse transcription Kit (Applied Biosystems, Waltham, MA, USA). The Primer-BLAST tool was used to design PCR primers (https://www.ncbi.nlm.nih.gov/tools/primer-blast/). RT-qPCR was performed in optical 96-well plates using SYBR® green in a LightCycler 480 (Roche, Basel, Switzerland). The *ubiquitin1* gene (*OsUbi1*, Os06g0681400) was used to normalize transcript levels. The 2^-ΔΔCt^ method was used to calculate relative expression levels. Four biological replicates each one from a pool of four different plants, and three technical replicates for each biological replicate were analysed. Primers used for RT-qPCR and genes analysed are listed in **Table S1**. Senescence-associated genes (SAGs) were selected from the Leaf Senescence Database 5.0 (LSD, https://ngdc.cncb.ac.cn/lsd/) (Zhao et al., 2024).

### 2.9. Statistical analysis

One-way ANOVA was used throughout the manuscript for Pi content, leaf senescence area, chlorophylls and carotenoids quantification, malondialdehyde content, electrolyte leakage, enzyme activity, and gene expression by qPCR. Two-way ANOVA was used for chlorophyll and carotenoid content analysis under methyl viologen treatment. One-and two-way ANOVA analysis was followed by Tukey’s multiple comparisons post hoc test (adjusted *P* ≤ 0.05). Data was analysed using GraphPad Prism v10.4.1 software (https://www.graphpad.com/) for statistical analysis.

## 3. Results

### 3.1. Pi treatment interferes with the photosynthetic pigment content and leaf senescence of rice plants

To investigate the adaptive responses of rice seedlings to Pi treatment, seeds were pre-germinated for 1 week in water and then transferred to soil. Seedlings were then supplied with varying concentrations of Pi, e.g., 0.025 mM Pi (hereinafter referred to as Low Pi plants), 1.0 mM Pi (Control Pi plants), and 2.5 mM Pi (High Pi plants) for 2 weeks, 3 weeks, or 4 weeks (**Figure S1A**). At 2 weeks of treatment, Pi levels progressively increased in leaves of rice seedlings (**Figure S1B**), indicating that rice plants sense and react to Pi treatment. Most importantly, after two weeks of Pi treatment, no significant differences were observed in growth of rice seedlings among conditions (**Figure S1C**). By extending Pi treatment, the seedlings continued accumulating Pi in their leaves (3 weeks and 4 weeks of treatment; **Figure S2A**, **B**). However, at 3 weeks or more of Pi treatment, seedling growth was significantly reduced in both Low-Pi and High-Pi plants compared with Control Pi plants (**Figure S2A**, **B**). Additionally, plants under high Pi developed leaf tip necrosis, which is a typical symptom of Pi toxicity in rice (**Figure S2C**). To avoid confounding effects caused by growth inhibition or Pi cytotoxicity, the short-term treatment (i.e., 2 weeks of Pi treatment) was adopted for studying adaptive responses to Pi in rice seedlings.

As previously mentioned, degradation of photosynthetic pigments, chlorophylls and carotenoids, occurs during leaf senescence causing leaf yellowing (Mayta et al., 2019). To note, HPLC analysis of leaf extracts revealed a progressive decrease in chlorophyll concentration, both chlorophyll *a* (Chl *a*) and chlorophyll *b* (Chl *b*), in leaves of the rice seedlings upon increasing Pi supply (**Figure 1A**). Chl *a* values ranged from 0.174 to 0.140 µg/mg dry weight in leaves of Low-Pi plants and High-Pi plants, respectively, while chlorophyll *b* values ranged from 0.062 to 0.048 µg/mg dry weight (Low Pi and High Pi plants) (**Figure 1A**). The positive impact of Pi on chlorophyl content raised the possibility of delayed senescence in low-Pi-treated rice seedlings. Similarly, a gradual reduction in carotenoids content, both carotenes (β-carotene, 9-cis-β-carotene) and xanthophylls (lutein, neoxanthin and violaxanthin) was observed when increasing Pi supply to the rice seedlings (**Figure 1B**).

**FIGURE 1.**
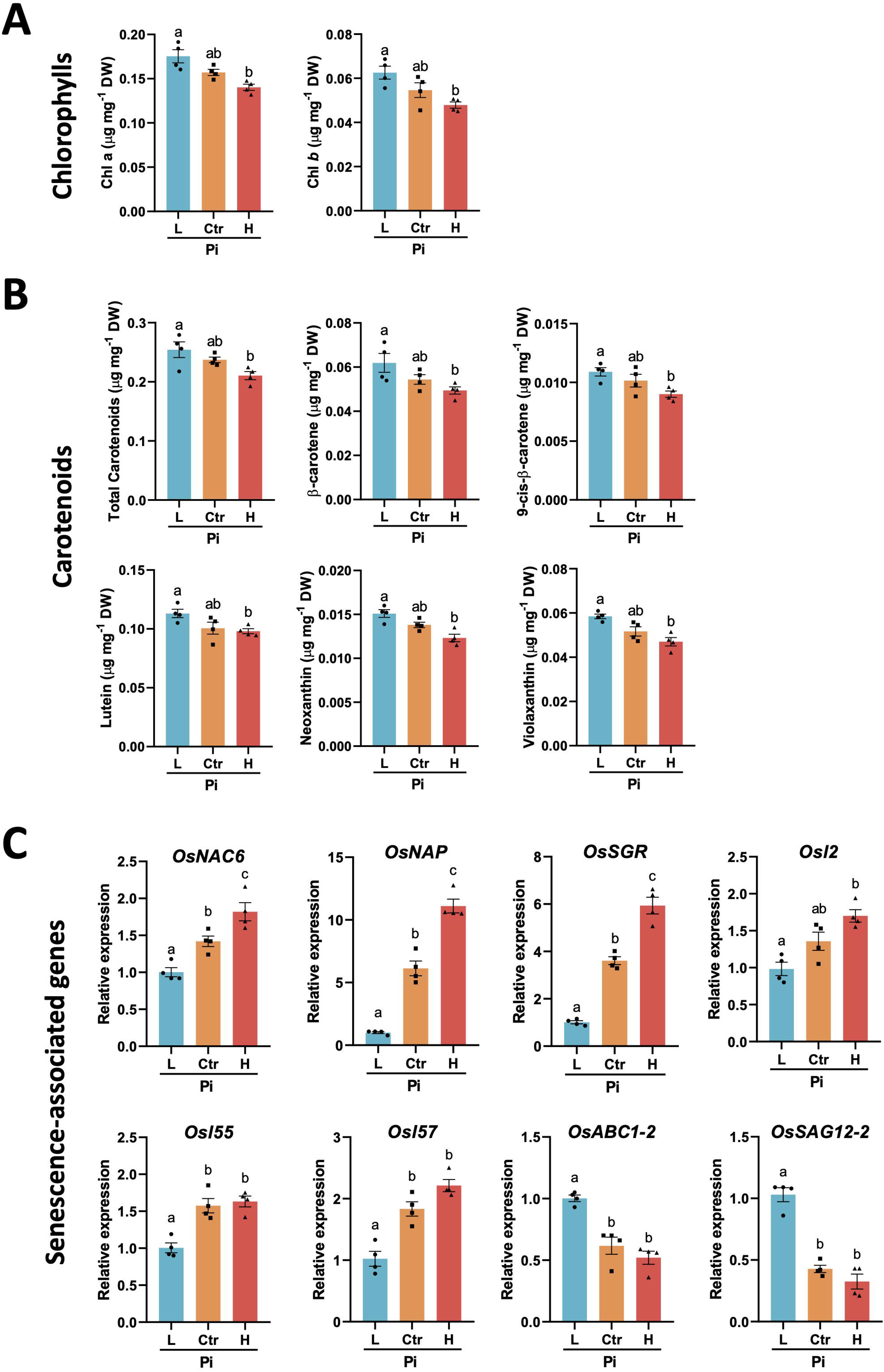
Effect of Pi treatment on chlorophyll and carotenoid content in rice leaves. Leaf extracts were prepared from seedlings that have been grown under different Pi supply (Low Pi, Control and High Pi seedlings) for 2 weeks. Chlorophyll and carotenoid content was determined by HPLC (A and B). Results shown correspond to the youngest fully developed leaves (Leaf 1; the same results were obtained in Leaf 2). (**A**) Chlorophyll content (Chlorophyll a, Chl *a*; Chlorophyll b, Chl *b*). (**B**) Content of total carotenoids, β-carotene, 9-cis-β-carotene, lutein, violaxanthin, and neoxanthin. Results are expressed as µg/mg of dry weight. (**C**) Expression of senescence-associated genes (SAGs): *OsNAC6*, *OsNAP*, *OsSGR*, *Osl2*, *Osl55*, *Osl57*, *OsABC1-2* and *OsSAG12-2*. Three independent experiments were conducted that produced the same results. Bars represent mean ± SEM of 4 biological replicates, each one from a pool of 4 different plants Statistically significant differences were determined by one-way ANOVA (different letters indicate significant differences among Pi conditions).

*SAG*s have a pivotal role in leaf senescence, some of them being directly involved in chlorophyll degradation (Guo et al., 2021). Knowing that short-term treatment of rice seedlings with low or high Pi can antagonistically delay or accelerate leaf senescence, respectively, it was of interest to investigate whether the expression of SAGs is affected by Pi treatment in rice seedlings. Accordingly, in this work, we monitored the expression of genes whose expression is known to be induced during senescence, such as *OsNAC6*, *OsNAP*, *OsSGR*, *Osl2*, *Osl55*, *Osl57* (Lee et al., 2001; Park et al., 2007; Liang et al., 2014; Wairich et al., 2023). Of them, *OsSGR* (*STAY-GREEN* encoding an enzyme involved in chlorophyll degradation during leaf senescence) is a marker for developmental leaf senescence (Park et al., 2007). In agreement with the above-described phenotype of senescence, all these SAGs exhibited a weaker expression in Low-Pi plants, their expression progressively increasing with increased Pi supply (**Figure 1C**). Conversely, *OsABC1* (encoding a kinase localized at the chloroplast envelope that helps to maintain chloroplast function during senescence) (Gao et al., 2012), and *OsSAG12* (a negative regulator of cell death, whose expression is induced during natural leaf aging) (Singh et al., 2016) showed higher expression in low-Pi relative to control and high-Pi rice seedlings (**Figure 1C**). In addition to *OsSGR*, several other genes involved in chlorophyll degradation (also often referred to as SAGs) also showed a weaker expression in low-Pi plants than in control and high-Pi plants (e.g., *OsNYC1*, *OsNOL*, *OsHCAR*, *OsNYC3*, *OsPAO* and *OsRCCR1*) (**Figure S3A**). Opposite pattern could be observed for genes required for chlorophyll biosynthesis, such as *OsGUN5*, *OsCRD1*, *OsPORA* and *OsPORB*, their expression being higher in low-Pi plants relative to control and high-Pi plants (**Figure S3B**).

Consistent with the observed reduction in carotenoid levels upon increasing Pi supply, transcript levels of central genes in the MEP (methylerythritol phosphate) and carotenoid biosynthetic pathways progressively decreased from low to high Pi conditions (**Figure S4**). They were: *OsDX3* (encoding the enzyme that catalyzes the initial step of the MEP pathway that produces precursors for carotenoids), *OsPSYI* (the central enzyme that determines the total amount of carotenoids accumulated in plant tissues), *OsPDS*, *OsLCYe* (carotenes biosynthetic pathway), *OsCYP97A4* and *OsHYD1* (xanthophylls biosynthetic pathway) (**Figure S4**).

Dark treatment has been shown to promote chlorophyll degradation and senescence (Sobieszczuk-Nowicka et al., 2018). Accordingly, the effect of Pi treatment on leaf senescence was further investigated using dark-induced leaf senescence (DILS) approaches. Initially, dark treatment was carried out at the whole plant level. For this, the rice plants grown under a regular photoperiod condition under different Pi regimes for 2 weeks, and 1 week more under continuous dark. Control plants were maintained under photoperiod condition over the entire period of time. Upon dark treatment, leaves of Low-Pi seedlings consistently showed less leaf yellowing than leaves from Control and High-Pi seedlings (**Figure 2A**). Thus, when increasing Pi supply, leaves showed a downward trend in chlorophyll content and carotenoid content (**Figure 2B, C**).

**FIGURE 2.**
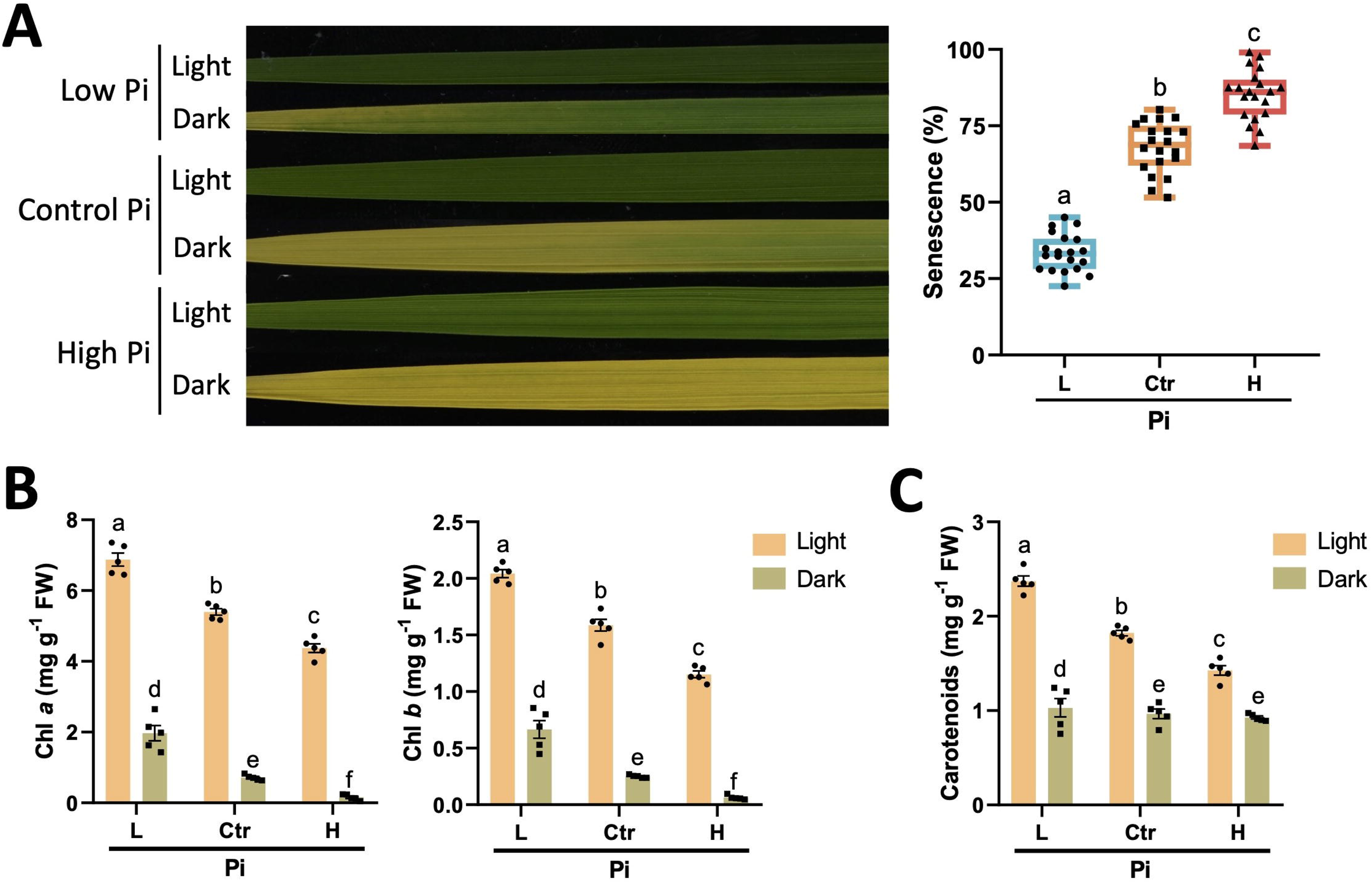
Dark-Induced Leaf Senescence (DILS) in rice seedlings supplied with increasing Pi concentrations. Whole leaves were detached from Low-Pi, Control Pi and High Pi) and subjected to dark treatment. (**A**) Representative images of leaves maintained in darkness for 9 days. Right panel, quantification of senescence regions at 9 days of dark treatment) by image analysis using the software APS Assess 2.0. The senescence level is represented as the % of the leaf area showing senescence relative to the total leaf area. Box plots represent median and data distribution (N = 20 leaves, each condition). (**B**) Chlorophyll content (Chlorophyll a, Chl *a*; Chlorophyll b, Chl *b*). (**C**) Total carotenoids content. Chlorophylls and carotenoids content (B and C) was determined by the spectrophotometric method and expressed as mg/g of fresh weight. Data shown correspond to Leaf 1 (similar results were obtained with Leaf 2). Bars in B and C represent mean ± SEM of 5 biological replicates, each one from a pool of 3 different plants. Statistically significant differences were determined by two-way ANOVA (different letters indicate significant differences among Pi conditions).

In rice, leaf senescence usually progresses from the leaf tip (consisting of the oldest cells) to the leaf base (youngest cells) (Gregersen et al., 2008). To follow leaf yellowing pattern with time along the leaf lamina, complementary DILS assays were conducted on entire leaves detached from Pi-treated rice seedlings. At 9 days of dark treatment, leaves from Low-Pi plants exhibited yellowing only at the leaf tip while the middle and basal regions of the leaf lamina stayed green (**Figure S5A**, left panel). By the same time, leaves from control and High-Pi plants showed severe yellowing along the entire lamina which increased when increasing Pi supply (**Figure S5A**, left panel). Colour image analysis confirmed delayed senescence in rice leaves of Low-Pi plants (25% vs 50-75% area of senescence) (**Figure S5A**, right panel). As it was observed in DILS assays with whole leaves, a slower progression of leaf yellowing over time was also observed in DILS assays with leaf sections from detached leaves from Low-Pi plants compared with leaves from Control and High Pi plants (**Figure S5B**). The Low-Pi leaves maintained green colour for a longer period of time. Furthermore, a gradual decline in chlorophyll content could be observed over time in leaf segments subjected to dark treatment, this reduction being more important when increasing Pi supply (**Figure S5C**). Equally, carotenoid content decreased when increasing Pi supply (**Figure S5D**).

Collectively, these analyses confirmed that Pi content in rice leaves modulates leaf senescence in rice seedlings. Short treatment with low Pi causes a delay in leaf senescence, while Pi accumulation accelerates this process.

### 3.2. Short-term treatment with low Pi decreases membrane lipid peroxidation and electrolyte leakage, and increases the antioxidant capacity of rice leaves

As our findings interwove Pi regimes with a senescent state, we further investigated the consequences of Pi treatment on senescence-related processes, particularly in processes related to ROS production. Under normal growth conditions, ROS accumulation during photosynthesis is controlled by antioxidant systems that prevent damage from oxidative stress. However, the over-generation of ROS can disrupt the equilibrium between ROS production and scavenging, which might cause membrane lipid peroxidation and eventually would affect membrane integrity and functionality. To address whether Pi treatment of rice seedlings has an effect on membrane lipid peroxidation, we determined the concentration of malondialdehyde (MDA) derivatives as a proxy of lipid peroxidation in cellular membranes. Complementary, electrolyte leakage measurements served to assess membrane permeability in leaves. Compared with Control Pi plants, the rice plants subjected to short-term treatment with Low-Pi plants accumulated significantly lower levels of MDA, these plants also showing lower electrolyte leakage values than control plants in their leaves (**Figure 3A**). Conversely, MDA concentration and electrolyte leakage values were higher in High-Pi plants than in Control Pi plants (**Figure 3A**). Our data pointed to higher membrane stability under short term low Pi treatment in rice plants.

**FIGURE 3.**
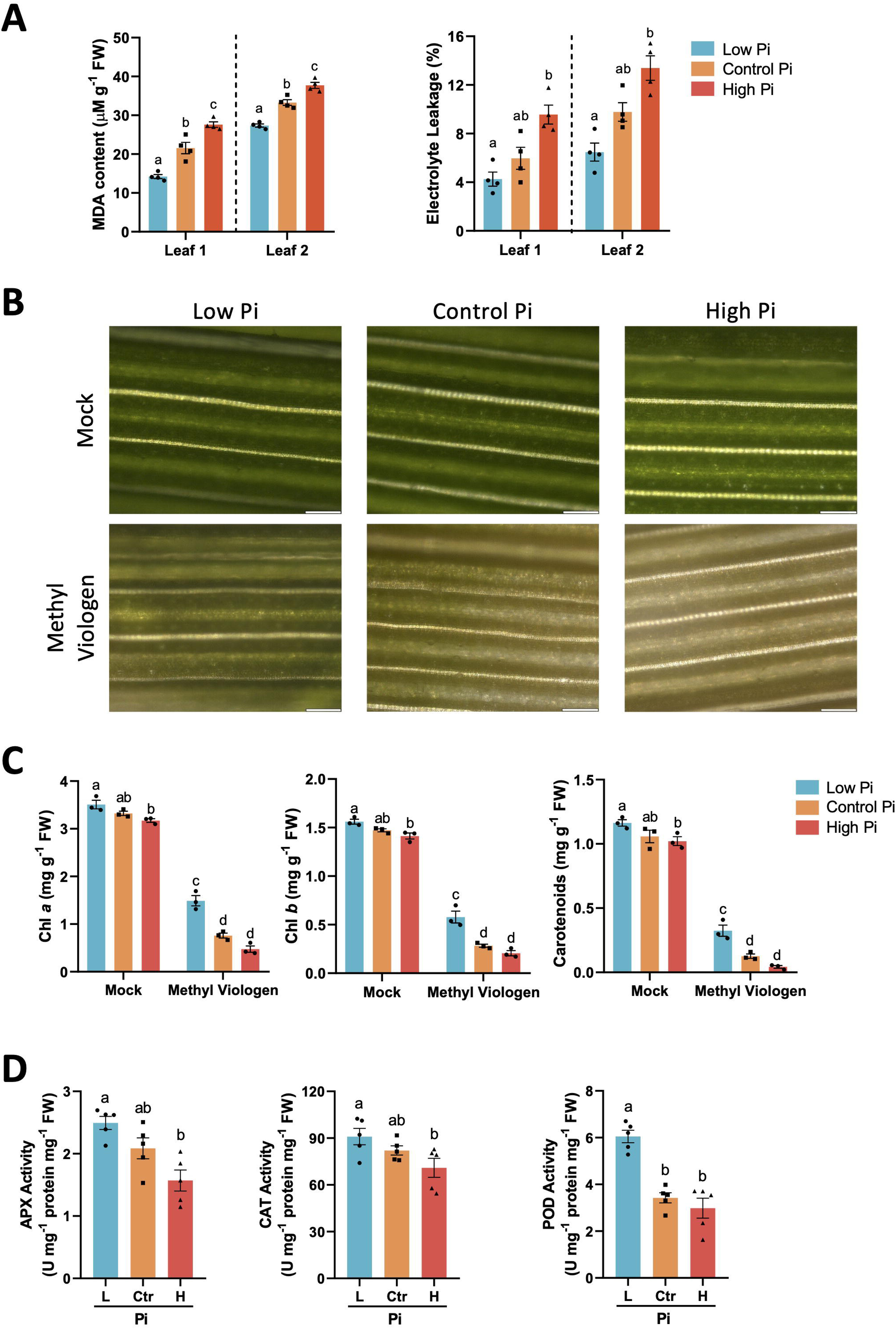
Effect of Pi treatment on senescence-associated processes. Pi-mediated alterations in membrane lipid peroxidation and permeability (A), tolerance to methyl viologen-induced oxidative stress (B, C), and antioxidant enzyme activities (D). Leaves from rice seedlings that have been grown under low Pi, Control Pi, or High Pi were examined. (**A**) MDA content (left panel) and electrolyte leakage (right panel). Leaf 1 and Leaf 2 from the same plant were examined separately. Four biological samples, each one from a pool of leaves from 6 different plants were analyzed. (**B**) Effect of methyl viologen (10 µM, MV) on rice leaves. Representative images of MV-treated leaves (Leaf 1) are shown (similar results were observed on Leaf 2). Water-treated leaves served as controls. Scale bars = 100 µm. (**C**) Quantification of photosynthetic pigments in MV-treated rice leaves. Chlorophylls (Chl *a*, Chl *b*) and carotenoids were extracted and quantified spectrophotometrically. Three biological replicates, each one from a pool of 4 different plants were examined. (**D**) Changes in the activity of ascorbate peroxidase (APX), catalase (CAT) and peroxidase (POD). Five biological replicates, each one from a pool of 6 different plants were examined. Three independent experiments were conducted with similar results. Bars in A, C and D, represent mean ± SEM. Statistically significant differences were determined by one-way ANOVA (**A**, **D**) or two-way ANOVA (**C**). The different letters indicate significant differences among Pi conditions.

Having found that the Pi nutritional state of the plant can alter the membrane lipid peroxidation and permeability we sought to evaluate tolerance to methyl viologen (MV)-mediated oxidative stress in leaves of Pi-treated plants. Exposure to MV (or paraquat) is known to induce ROS production in chloroplasts (Scarpeci et al., 2008). Compared with Control Pi plants, MV-induced chlorophyll bleaching was less pronounced in plants that have been treated with Low-Pi, while leaves of High-Pi plants were more sensitive to MV-induced oxidative stress (**Figure 3B**). In line with visual inspection of MV-treated leaves, chlorophyll and carotenoid content was significantly higher in MV-treated leaves of Low-Pi plants than in leaves of Control and High-Pi plants (**Figure 3C**).

It is also known that, if the ROS molecule H_2_O_2_ that is generated during photosynthesis is not adequately eliminated, its accumulation leads to the formation of hydroxyl radicals, further contributing to oxidative stress. The enzymes Ascorbate Peroxidase (APX), Catalase (CAT) and Peroxidase (POD) are responsible for detoxifying H_2_O_2_. Interestingly, the activity of APX, CAT and POD was significantly higher in leaves of Low-Pi plants compared to Control and High-Pi plants (**Figure 3D**). The observed tolerance to MV-induced oxidative stress in leaves of Low-Pi seedlings might well be the consequence of higher levels of antioxidant enzyme activities (e.g., APX, CAT and POD), which is expected to hamper H_2_O_2_ accumulation in chloroplasts. In this way, short-term treatment with low Pi appears to exert a protective effect against oxidative stress in rice leaves.

### 3.3. Transcriptome profiling reveals an orchestrated reconfiguration of central biological processes in leaves of rice plants treated with low Pi

To obtain a comprehensive view of the molecular mechanisms underlying the adaptive response of rice plants to treatment with low Pi, we conducted RNA-Seq profiling of leaves from Low-Pi and Control Pi plants (**Table S2**). Principal component analysis (PCA) clearly separated transcriptome responses under each Pi treatment (93% variance in PC1) (**Figure S6A**). Differentially expressed genes (DEGs) in leaves of plants under low Pi were declared based on absolute log_2_ fold change (FC) ≥ 0.5 to Control Pi (adjusted *P* ≤ 0.01, Wald’s test). Hierarchical clustering analysis of the DEGs further illustrated that differences in transcriptomes between low Pi-treated and control plants covered transcripts with different steady-states (**Figure S6B**). A similar number of DEGs was found to be up-regulated and down-regulated, namely 2908 up- and 2945 down-regulated, respectively. The complete list of DEGs is presented in **Table S3**.

To functionally analyse the relevance of DEGs, significant enrichment in non-redundant Gene Ontology (GO) terms was conducted for biological processes and summarised using REVIGO (Supek et al., 2011). GO terms with highest fold enrichment were related to “Metabolic processes” for both up- and down-regulated genes (**Figure 4A**; **Table S4**). Notably, “Photosynthesis and Light harvesting”, “Generation of precursor metabolites and energy”, and “Pigment biosynthetic processes” were over-represented in the set of up-regulated genes in Low-Pi plants (**Figure 4A**, left panel). A similar trend was observed for terms related to “Response to stimulus” (e.g., “Cellular response to phosphate starvation”, “Response to nutrient levels”, “Cellular detoxification”), “Ion transport” (e.g., “Phosphate ion transport”), “Plant-type cell wall biogenesis” and “Sulfolipid biosynthetic process” (**Figure 4A**, left panel; **Table S4**). Conversely, “Metabolic processes”-related terms for down-regulated genes in Low-Pi plants were mainly linked to “Nitrate assimilation” (**Figure 4A**, right panel; **Table S4**).

**FIGURE 4.**
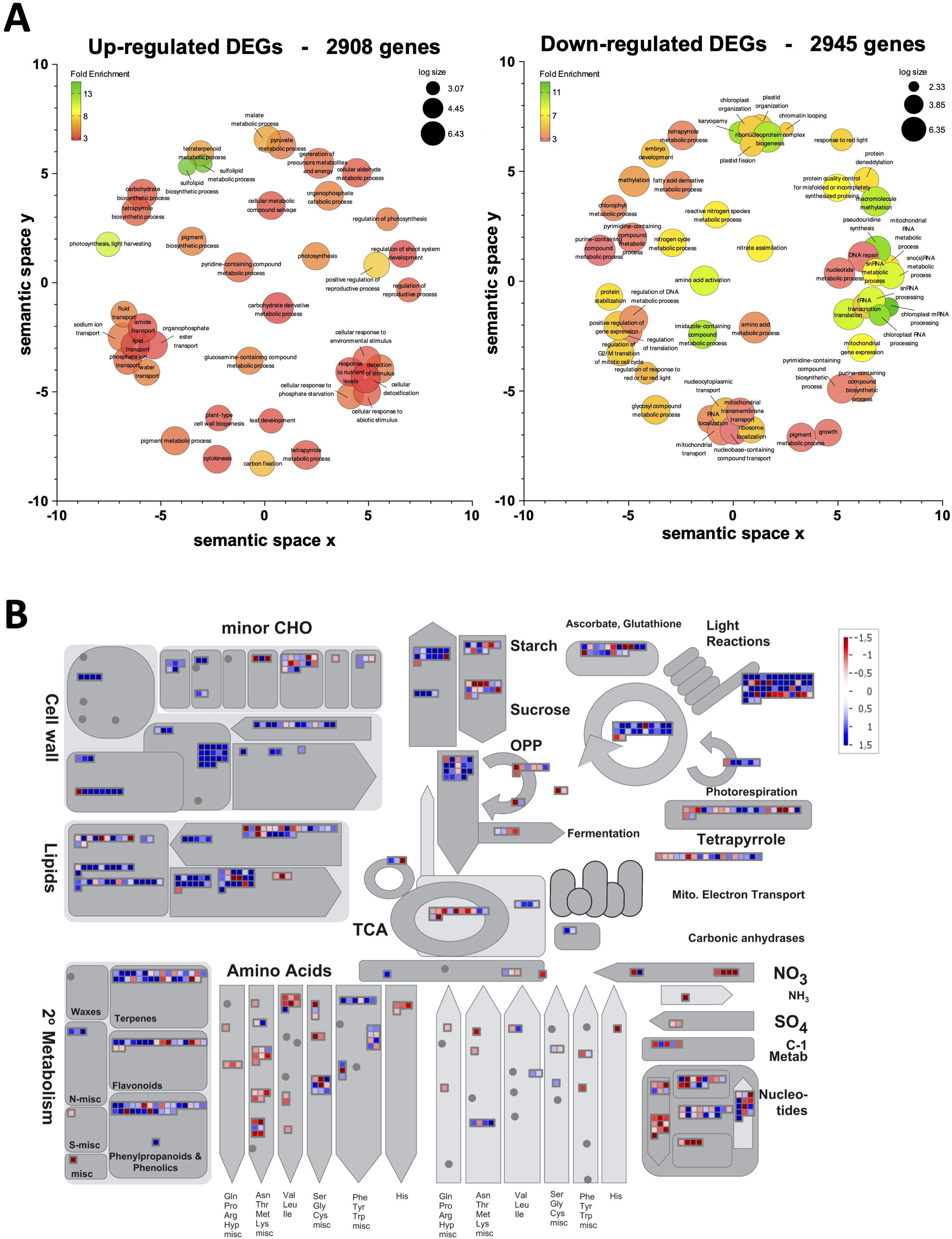
Analysis of genes that regulated by treatment with low Pi in leaves of rice plants. (**A**) Gene Ontology (GO) terms significantly enriched in leaves of rice plants treated with Low-Pi. GO terms enriched in the set of up-regulated genes (left panel) and down-regulated genes (right panel). The most enriched GO terms (number of DEGs ≥ 5; Enrichment score ≥ 2.5; number of genes < 900) were considered and visualized using REVIGO after reducing redundancy and clustering of similar GO terms. Disc colour (blue to red) indicate the fold enrichment and disc size is proportional to the frequency in the GO *Oryza sativa* database (e.g., larger and smaller discs represent more general and more specific GO terms, respectively). Full list of genes in enriched GO terms is presented in Supplemental Table 2). (**B**) MapMan overview of plant metabolism. Individual genes are represented by small squares. The colour key represents the log2FC (values scaled to 1.5 to −1.5). Red represents the down-regulation and blue represents the upregulation of rice genes in leaves of Low-Pi rice plants relative to control plants.

Since central metabolism was importantly targeted by Low-Pi treatments, a MapMan classification of DEGs was used to dissect the altered pathways in plant metabolism. Low-Pi led to an overall increase in levels of most of the depicted pathways, namely carbon-related metabolism (minor carbohydrates, lipids, starch/sucrose, Calvin cycle, oxidative pentose pathway) (**Figure 4B**; **Figure S7**). On the other hand, Low-Pi treated plants displayed lower levels for transcripts related to amino acid and nucleotide catabolism (**Figure 4B**; **Figure S7**). This result tempted us to propose that plants under low Pi might be more efficient in autotrophy (light reactions and production of carbon skeletons) and less dependent on heterotrophy (organic acid and protein catabolism, nucleic acid recycling).

The higher autotrophic capacity of Low-Pi plants is consistent with the increased concentration of photosynthetic pigments (chlorophylls and carotenoids) and the delayed senescence observed under this condition (see **Figure 1** and **Figure 2**). The RNA-Seq datasets were further explored to retrieve genes involved in metabolism of photosynthetic pigments, as well as genes involved in chloroplast biogenesis and development (e.g., *OsPSYI*, *OsPDS*, *OsHYD1*, *OsNOL*, *OsNYC3*, *OsPAO*, *OsRCCR1*, *OsGUN5*, *OsCRD1*, *OsPORA*, *OsPORB*) (**Table S5**). In the same line, the expression of an important number of *SAG* genes (e.g., *OsNAC6*, *OsNAP*, *OsSGR*, *Osl2*, *Osl55*, *Osl57*, *OsABC1-2*, *OsSAG12*-*2*) and regulatory pathways associated with leaf senescence were regulated in Low-Pi plants (**Table S6**). Finally, consistent with results obtained on tolerance to MV-mediated oxidative stress and antioxidant activities, transcripts encoding enzymes involved in redox-homeostasis (e.g., peroxidases, ascorbate peroxidases, thioredoxins, glutathione-S-transferases, glutaredoxins and catalases, among others) were also found to be regulated in rice plants under low Pi treatment (**Table S7**). Collectively, our transcriptome data provided a broad picture of the molecular regulatory mechanisms involved in the rice response to short-term treatment with low Pi. Transcriptional alterations in gene expression were significantly enriched in a vast number of metabolic pathways and biological processes representing adaptive responses to short-term treatment with low Pi in rice rather than a typical general stress response.

In Molecular Function, the most abundant terms in the set of up-regulated genes were “Glycosyltransferase activity”, “Phosphoric ester hydrolase activity” and “UDP-glycosyltransferase activity”, while down-regulated genes classified in diverse molecular functions, including “Catalytic activity on RNA” and “Hydrolase activity” (**Table S8**). In the category of Cellular Components, the top four significantly enriched terms among up-regulated genes were “Plastid envelope”, “Thylakoid”, “Outer Membrane” and “Photosynthetic Membrane” (**Table S9**). Further interpretation of biochemical functions of the DEGs in Low Pi plants came from PANTHER-PC ontology (Protein ANalysis THrough Evolutionary Relationships-Protein Class) analyses (Mi et al., 2021). This analysis revealed that the Protein Class of “Hydrolase” was the most frequent class in the set of up-regulated genes in Low-Pi plants, which included enzymes that are predominantly associated with Pi remobilization (e.g., phosphosphodiesterases, Phosphatases, and Phospholipases, among others) (**Table S10**). Important key genes related to adaptation to Pi deficiency that were also regulated in Pi-treated plants, including Pi transporter genes, the Pi-inducible *OsPHR4* transcription factor, *SPX domain-containing protein* genes, *IPS1* and *IPS2*, as well as genes encoding ribonucleases (S*-RNases* or *RNSs, PR10a* or *PBZ1*) and genes required for the biosynthesis of non-phosphorus lipids, both galactolipids and sulfolipids (**Table S11**). In summary, our transcriptomic analysis revealed a mechanism of global gene expression regulation triggered by short-term treatment with low Pi in rice. Diverse metabolic pathways and physiological functions contributing to delayed leaf senescence seen to be regulated in the plant’s adaptive response to treatment with low Pi.

### 3.4. Alterations in the expression of *MIR827* and *MIR399* has an effect on leaf senescence in rice

As a complementary approach to Pi treatment of rice plants, we investigated the impact of overexpression of *MIR827* or *MIR399* on senescence-associated mechanisms. Previous studies revealed that rice plants constitutively expressing *MIR399* or *MIR827* is accompanied by an increase in Pi content in rice leaves (Campos-Soriano et al., 2020; Bundó et al., 2024). Conversely, silencing *MIR827* by CRISPR/Cas9-mediated genome editing decreases Pi content (Bundó et al., 2024).

Homozygous miR827 overexpressor lines (miR827 OE; 3 independently generated lines) and CRISPR/Cas9-edited rice lines harbouring two different mutations in the *MIR827* locus (CRISPR-miR827 lines) were assayed (Bundó et al., 2024). As previously reported, these plants did not display obvious phenotypic differences relative to wild-type at the three-leaf stage (**Figure S8A**, **B**, left panels). At a later stage developmental stage, however, miR827 OE plants showed leaf tip necrosis caused by Pi overaccumulation as it was also described for miR399 OE plants (Campos-Soriano et al., 2020; Bundó et al., 2024). Compared with wild-type plants, miR827 OE and CRISPR-miR827 plants showed an increase or a decrease in Pi levels, respectively (**Figure S8A**, **B**, right panels). Notably, miR827 OE plants consistently showed accelerated leaf yellowing, whereas CRISPR-miR827 lines exhibited delayed senescence (**Figure 5A**; **Figure S9**). Moreover, leaves of miR827 OE plants were completely bleached upon MV treatment, while MV-treated leaves of CRISPR-miR827 plants stayed greener than MV-treated leaves from wild-type plants (**Figure 5B**, **C**, upper panels). Consistently, the content of photosynthetic pigments in MV-treated leaves was reduced in miR827 OE relative to MV-treated wild-type plants (**Figure 5B**, lower panels). Conversely, photosynthetic pigments content was higher in MV-treated CRISPR-miR827 leaves than in MV-treated wild-type leaves (**Figure 5C**, lower panels).

**FIGURE 5.**
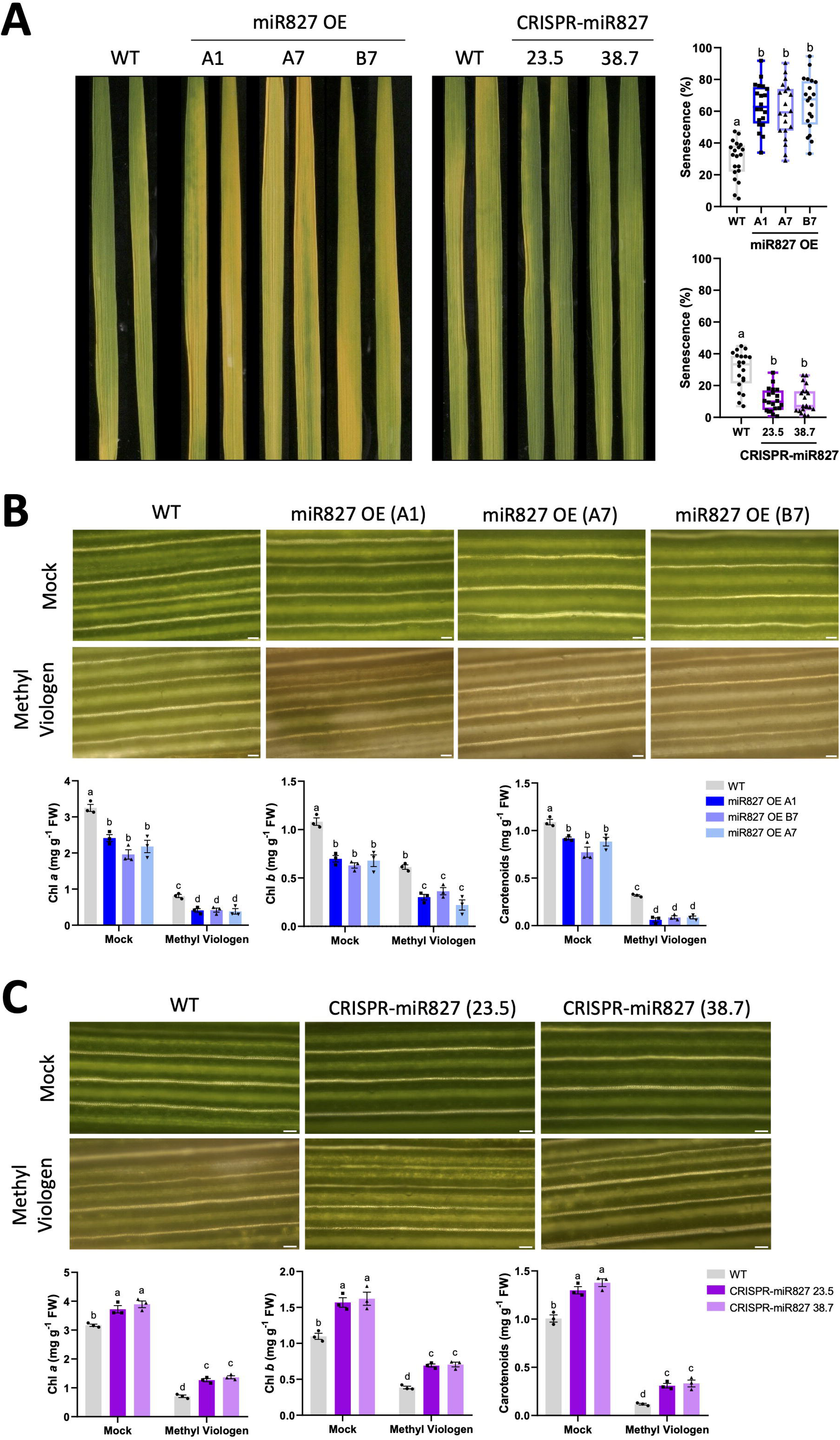
Analysis of senescence-associated processes in rice lines with altered expression **of *MIR827***. DILS assays were carried out on entire leaves (Leaf 1) collected from miR827 overexpressor (miR827 OE) and CRISPR/Cas9-edited plants (CRISPR-miR827) plants at the 3-4 leaf stage. Leaves were maintained in darkness for 6 days in the case of the miR827 OE lines, or 9 days in the case of the CRISPR-miR827 lines. (**A**) Representative images of senescing leaves (entire leaves) subjected to DILS (left panel). Right panel, quantification of senescing regions. The senescence level is represented as the % of the leaf area showing senescence relative to the total leaf area. Box plots represent median and data distribution (N = 20 leaves, each genotype). Image analysis was carried out using the software APS Assess 2.0. (**B**) Effect of methyl viologen (MV) treatment on leaves (Leaf 1) from miR827 OE rice plants. Upper panels, representative images of leaf sections treated with MV or water-treated leaves (mock). Bars = 50 µm. Lower panels, spectrophotometric measurement and quantification of photosynthetic pigments in MV-treated rice leaves, chlorophylls (Chl *a*, Chl *b*) and carotenoids. Bars represent mean ± SEM of 3 biological replicates, each one from a pool of 4 different plants. (**C**) Effect of methyl viologen (MV) on leaves (Leaf 1) of CRISPR-miR827 rice plants. Upper panels, representative images of leaf sections treated with MV (10 µM) or water-treated leaves (mock). Scale bars = 50 µm. Lower panels, spectrophotometric measurement of photosynthetic pigments in MV-treated rice leaves, chlorophylls (Chl *a*, Chl *b*) and carotenoids. Bars represent mean ± SEM of 3 biological replicates, each one from a pool of 4 different plants. Two independent experiments were conducted with similar results. Statistically significant differences were determined by one-way ANOVA (A) and two-way ANOVA (B and C) (different letters indicate significant differences).

As previously noted in wild-type plants treated with high Pi, chlorophyll content (Chl *a* and Chl *b*) decreased in miR827 OE plants relative to wild-type plants, while their levels increased in CRISPR-miR827 plants (**Figure 6A**). Regarding carotenoids, their levels were lower in miR827 OE plants than in wild-type plants, but higher in CRISPR-miR827 plants (**Figure 6B**). As also noted in miR827 OE lines, the miR399 OE plants exhibited accelerated leaf senescence (**Figure S10A**, **B**), higher sensitivity to MV-induced oxidative stress (**Figure S10C**), and reduced levels of chlorophylls and carotenoids (**Figure S10D** and **Fig. S11**). Thus, results observed in miR399 OE and miR827 OE plants, both of them accumulating Pi in their leaves, correlated well with those obtained on wild-type plants that have been supplied with high-Pi. These findings reinforce the notion that Pi content in rice leaves regulates leaf senescence.

**FIGURE 6.**
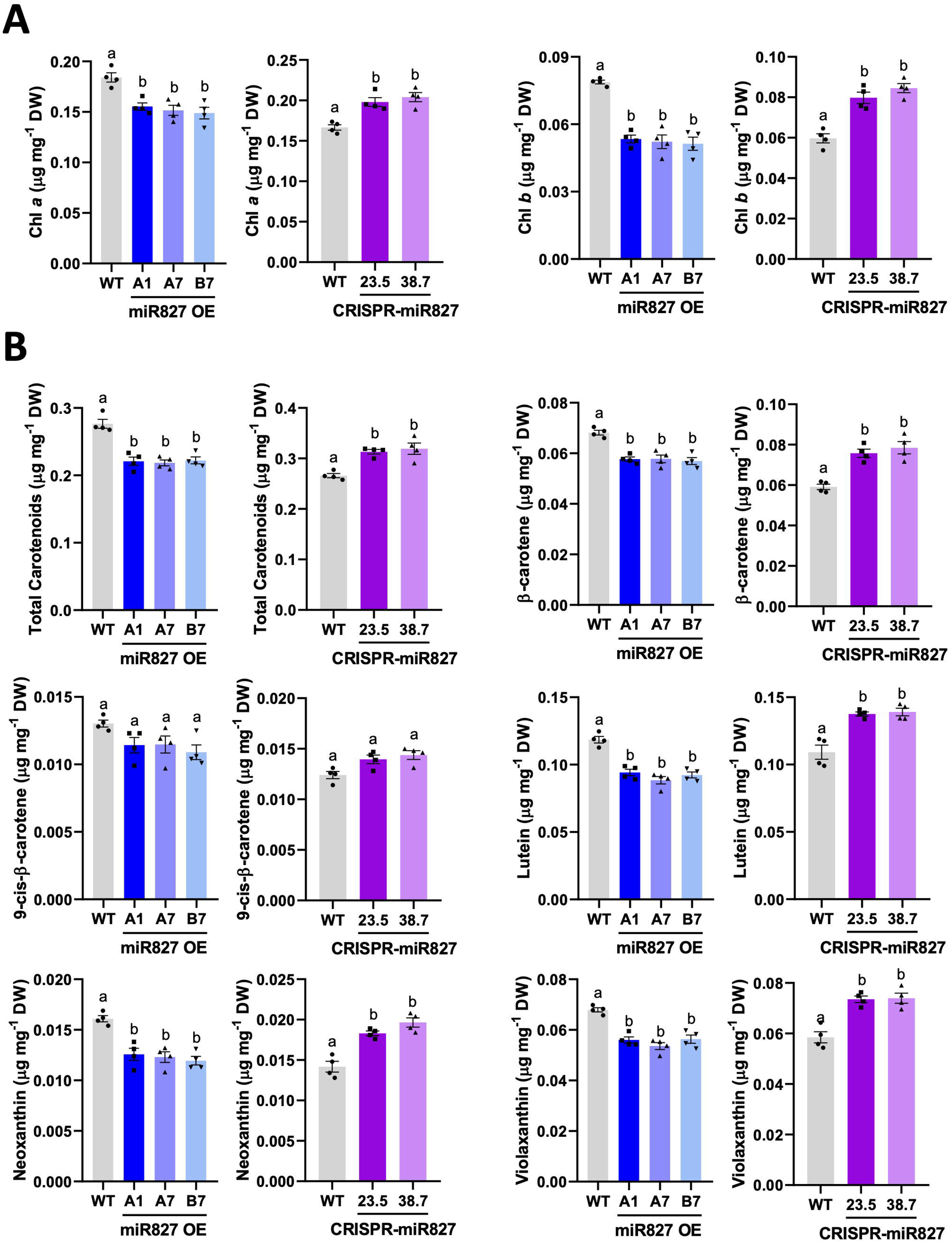
Content of photosynthetic pigments in leaves of rice plants with altered expression of *MIR827*. Leaves were collected from miR827 overexpressor (miR827 OE) and CRISPR/Cas9-edited plants (CRISPR-miR827) at the 3-4 leaf stage (Leaf 1, youngest leaf). Chlorophylls and carotenoids content was determined by HPLC. (**A**) Chlorophyll *a* and Chlorophyll *b* content. (**B**) Carotenoid content (total carotenoids, lutein, 9-cis-β-carotene, β-carotene, violaxanthin and neoxanthin). Bars represent mean ± SEM of 4 biological replicates, each one from a pool of 4 different plants. Three independent experiments were conducted. Statistically significant differences were determined by one-way ANOVA (different letters indicate significant differences among Pi conditions).

## 4. Discussion

In this study we provide evidence that Pi has a strong impact on leaf senescence during early vegetative growth in rice. Supporting this conclusion, qualitative and quantitative analyses of senescence-related parameters on rice seedlings under different Pi nutritional regimes demonstrated that short-term Pi treatment delays leaf senescence in young rice plants while increasing Pi supply triggers a senescent leaf state. Not only the Pi fertilization regime but also alterations in MIR827 and MIR399 expression alter the program of leaf senescence. Decreasing Pi content in CRISPR-Cas9-edited *MIR827* rice seedlings causes a delay in leaf senescence, while increasing Pi content in *MIR399* and *MIR827* plants accelerates leaf senescence. Importantly, in this study, short-term treatments were applied to rice seedlings to avoid negative effects on plant development observed upon longer periods of Pi treatment (e.g., reduction in plant growth in Low-Pi plants or Pi toxicity in High-Pi plants), an aspect that has not been accurately reflected in other studies. Clearly, the strength (e.g., Pi concentration used) and duration of Pi treatment might determine how the plant responds to treatment, hence, how Pi content might influence senescence. Furthermore, this study was carried out in rice plants at an early developmental stage to avoid age-dependent factors controlling the initiation and progression of leaf senescence. Under these experimental conditions, confounding effects due to nutrient remobilization from aging leaves to younger, developing, or reproductive parts of the plant typically occurring in adult plants can also be ruled out. Alterations in the rate of leaf senescence observed in Pi-treated rice seedlings can then be attributed to Pi supply. The observation that Low-Pi treatment delays leaf senescence was an unexpected finding because nutrient deficiency can be expected to trigger senescence to facilitate nutrient remobilization to developing and reproductive tissues (Zhang et al., 2024a). This phenomenon has been mainly described in adult plants (age-dependent regulation of senescence), and/or plants under continuous Pi deprivation, a situation that severely interferes with physiological processes essential for plant growth, development.

Degradation of photosynthetic pigments is the most obvious hallmark of leaf senescence. Along with this, chlorophyll and carotenoid content was found to be higher under low-Pi availability, their content progressively decreasing when increasing Pi supply. A higher accumulation of photosynthetic pigments would then explain delayed senescence in leaves subjected to short-term low Pi treatment. These results align with those recently reported on an *indica* rice variety (e.g., KDML105 variety) using hyperspectral imaging, where Pi deficiency was found to increase chlorophyll content in Pi-deficient rice seedlings (Jaisue et al., 2025). Low Pi-induced delayed leaf senescence might well constitute an adaptive mechanism to sustain plant growth under temporary drops in P bioavailability in soils. Conversely, our results indicated that high Pi causes a significant reduction in chlorophyll content and accelerates leaf senescence, pointing to excess Pi being a stressful factor. Collectively, our results provide evidence towards a more resilient state in rice plants under limiting Pi, while Pi accumulation would hamper the plant capacity to cope with a stressful situation.

Results here presented support that short treatment of rice plants with low Pi confers tolerance to MV-induced ROS production in chloroplasts. An efficient antioxidant system (e.g., H_2_O_2_ detoxifying enzymes) in Low-Pi plants would explain, at least in part, the observed phenotype of tolerance to MV-induced oxidative stress in these plants. Opposite to this, Pi accumulation in leaves enhances sensitivity to MV-induced oxidative stress while decreasing antioxidant enzyme activities. A weakened capacity in scavenging ROS in High-Pi plants would make Pi-accumulating plants unable to properly cope with MV-induced oxidative stress. This fact is also in tune with higher lipid peroxidation and membrane permeability in High-Pi plants which are inherent features during leaf senescence. Additionally, the observed increase in carotenoid content in leaves of rice plants under low Pi supply is expected to protect membrane systems from lipid peroxidation and electrolyte leakage (Jaisue et al., 2025). Supporting this possibility, carotenoids are known to function as ROS scavengers to protect against oxidative damage in chloroplast, particularly the single oxygen (^1^O_2_), a highly ROS molecule produced in chloroplast, and the main cause of lipid peroxidation and chlorophyll bleaching (Falk and Munné-Bosch, 2010). Lower concentrations of carotenoids in leaves of High-Pi seedlings would then favour membrane lipid peroxidation and electrolyte leakage. Again, short treatment with low Pi appears to enhance the plant’s capability to withstand oxidative stress, whereas Pi accumulation represents a stressful factor negatively affecting ROS homeostasis in rice leaves.

Expression analysis of Pi-deprived rice plants provided a global view of the molecular framework underlying low-Pi-induced delayed leaf senescence in rice. In agreement with the observed phenotype of leaf senescence, low Pi treatment was accompanied by alterations in the expression of SAGs and genes required for the biosynthesis and/or degradation of photosynthetic pigments, both chlorophylls and carotenoids. Transcriptome analysis uncovered a molecular framework illustrating adaptive responses to short treatment with low Pi in rice, a process in which multiple biological processes appear to operate to effectively delay leaf senescence. For instance, among up-regulated genes in Low-Pi plants, there were genes involved in the biosynthesis of non-phosphorus lipids, both galactolipids (monogalactosyldiacylglycerol synthases and digalactosyldiacylglycerol synthases), and genes involved in the biosynthesis of sulfolipids (UDP-sulfoquinovose synthase; Sulfoquinovosyl transferase). Galactolipids in chloroplasts are essential for maintaining a functional photosynthetic apparatus. Under stress conditions, however, membrane phospholipids are substituted by non-phosphorus lipids, a situation in which galactolipids serve as a substitute for phospholipids in extraplastidic membranes (Okazaki et al., 2013; Li and Yu, 2018). Presumably, membrane lipid remodelling switching from phospholipids to galactolipids and sulfolipids is expected to occur in cellular membranes of Low-Pi rice plants. Transcript profiling also revealed that Low-Pi plants display a more efficient autotrophic state, as genes involved in carbon-related pathways, secondary metabolites and lipid metabolism are expressed at higher levels in these plants. Genes whose expression is regulated by short treatment with low Pi treatment in rice seedlings can then be used as functional markers to build up strategies based on Pi treatments aiming to delay leaf senescence in rice.

In addition to treatment with low Pi, alterations in *MIR827* or *MIR399* expression has been shown to have an impact on leaf senescence in rice. As previously mentioned, CRISPR/Cas9-mediated mutagenesis of *MIR827* decreases Pi content, whereas overexpression of one or another of these *MIR* genes, *MIR827* and *MIR399,* causes an increase in Pi levels in rice leaves (Campos-Soriano et al., 2020; Bundó et al., 2024). CRISPR/Cas9-mediated silencing of *MIR827* expression is accompanied by an increase in photosynthetic pigments content, delayed leaf senescence, and tolerance to MV-induced oxidative stress. Conversely, miR827 overexpression and miR399 overexpression (Pi-accumulating plants) causes a reduction in photosynthetic pigments and acceleration of leaf senescence and increases sensitivity to MV-induced oxidative stress in chloroplasts. In this way, both strategies, treatment of wild-type rice plants with high Pi and *MIR827* (or *MIR399*) overexpression, have similar consequences in processes related to leaf senescence. Furthermore, these results highlight the relevance of miR827 and miR399 in controlling both Pi homeostasis and leaf senescence in rice, further supporting the existence of shared regulatory mechanisms in Pi homeostatic and senescence programs. Senescence-associated processes are regulated at multiple levels, and miR399 and miR827 might well function in signal integration between Pi signalling pathways and senescence signalling pathways to rapidly adapt to changing nutrient environments.

Considering changes in the temporal pattern of leaf senescence, phenotype, expression of SAGs and other senescence-related physiological parameters, ROS detoxification, and genes controlling metabolic processes, and Pi levels, it can be concluded that multiple processes are involved in the adaptive response of rice to short treatment with low Pi. A model summarizing these adaptive responses to short Pi treatment is presented in **Figure 7**. Briefly, short term treatment with low Pi (as well as *MIR827* silencing), and subsequent decrease in Pi level, increases photosynthetic pigments content, chlorophylls and carotenoids, and enhances the activity of H_2_O_2_ detoxifying enzymes, which together with a higher carotenoids content would contribute to maintenance of ROS homeostasis and reduced levels of membrane lipid peroxidation and maintenance of membrane permeability. Treatment with low Pi is also accompanied by alterations in metabolic processes (e.g., those related to carbon metabolism, lipid metabolism or secondary metabolism which, in turn, might be responsible of delayed leaf senescence in low-Pi-treated plants. The opposite effects are observed in leaves of rice plants accumulating Pi either by fertilization with high Pi or by *MIR827* or *MIR399* overexpression.

**FIGURE 7.**
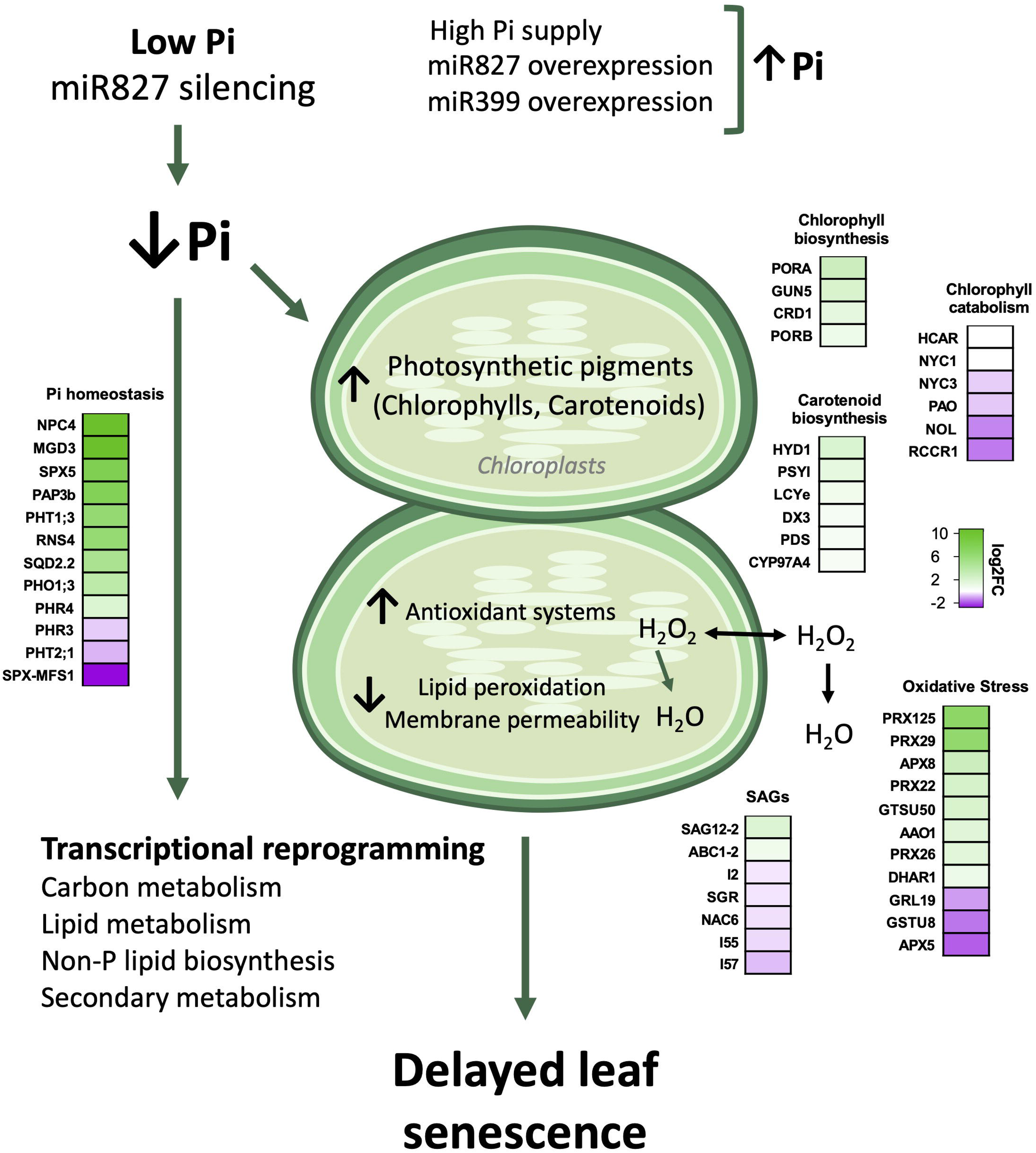
Mechanisms involved in Pi-mediated regulation of leaf senescence in rice plants. A decrease in Pi content caused by low Pi fertilization (or *MIR827* silencing) triggers an increase in photosynthetic pigments content, a reduction in membrane lipid peroxidation and permeability, and confers tolerance to MV-induced oxidative stress in chloroplast which is also associated with increased activity of enzymes involved in ROS detoxification. Decreasing Pi content in rice leaves also leads to alterations in the expression of genes involved in senescence-associated processes (e.g., SAGs, chlorophyll biosynthesis and degradation, carotenoid biosynthesis, oxidative stress), and genes involved in Pi homeostasis. An opposite response occurs in rice plants accumulating Pi (e.g., upon fertilization with high Pi or overexpression of miR827 and miR399) in which leaf senescence is accelerated compared with control plants. Notably, treatment with low Pi also regulates the expression of genes involved in diverse metabolic and physiological processes, such as carbon metabolism, lipid metabolism and secondary metabolism. Thus, this study uncovered a molecular framework illustrating mechanisms underlying low-Pi-induced delayed leaf senescence in rice.

Collectively, these results support that Pi nutrition is crucial for the control of leaf senescence in rice. Despite rice being one of the most important staple food, rice cultivation faces multifaceted challenges, including climate change, impact of pests and diseases, water eutrophication, primarily through excessive use of fertilizers, and anthropogenic pressures linked to urban expansion and concomitant loss of agricultural land. Deciphering processes by which rice plants adapt to changing Pi levels and mechanisms contributing to delayed leaf senescence is then of great importance in rice cultivation. On the other hand, the adoption of Green Revolution strategies during the last century greatly increased rice yields in part through the widespread use of fertilizers, which has led to a situation of Pi excess in rice fields. Therefore, modern rice varieties are highly dependent on Pi fertilizer application. Re-assessing current practices in rice cultivation based on the over-use of Pi fertilizers is then a need. Results here presented provide a theoretical basis to understand adaptation strategies to low Pi supply in rice that might help in reducing environmental problems caused by excess fertilization.

## Acknowledgements

We thank Dr Beatriz Val-Torregrosa, Glòria Escolà, Laia Castillo for assistance in sample processing and Iratxe Busturia for assistance in enzyme activity assays. We also thank Plant Growth and Metabolomics facilities at CRAG. This research was supported by project PID2021-128825OB-I00 (B. SS) and PID2024-162615OB-I00 (A.G-M) funded by MICIU/AEI/10.13039/501100011033 and by ERDF/EU; and grant RYC2022-037020-I (A. G-M) funded by MICIU/AEI/10.13039/501100011033 and by ESF+. We also acknowledge financial support from CEX2019-000902-S funded by MICIU/AEI/10.13039/50110001103 and the CERCA Program/Generalitat de Catalunya. H. M-C was a recipient of a fellowship from the MCIN (ref PRE2019-087477).

## Author Contribution

B.SS conceived the project H. M-C, M. B. and B.SS designed the experiments and analysed the data; H. M-C and M. B. performed the experiments; A. G-M analysed and discussed data on RNASeq. B.SS and H. M-C wrote the paper. All authors approved the final manuscript.

## Data availability Statement

The RNA sequence datasets generated in this study can be found at the National Center for Biotechnology Information (NCBI) Gene Expression Ommibus (GEO) with the accession number GSE298198 (GSE298198 (token, cbunqkssvpwpvip).

## Supporting Information

**Figure S1:** Conditions used for Pi treatment of rice seedlings. **Figure S2:** Effect of treatment of rice seedlings with Pi for 3 and 4 weeks. **Figure S3:** Expression of genes involved in chlorophyll metabolism in leaves of rice plants that have been grown under different Pi conditions. **Figure S4:** Expression of genes involved in carotenoid biosynthesis in rice plants grown under different Pi supply. **Figure S5:** Senescence phenotype of detached leaves from rice seedlings grown under increasing Pi supply for 2 weeks. **Figure S6:** Analysis of RNASeq data from leaves of Low-Pi and Control plants. **Figure S7:** MapMan overview of Photosynthesis. **Figure S8:** Phenotype of miR827 overexpressor and CRISPR/Cas9-edited miR827 rice plants. **Figure S9:** Senescence phenotype of detached leaf sections from miR827 OE or CRISPR-miR827 plants subjected to darkness during 1, 3 and 6 days. **Figure S10:** Senescence phenotype of leaves from rice plants overexpressing miR399. **Figure S11**: Content of photosynthetic pigments, chlorophylls and carotenoids, in leaves of miR399 OE rice plants. **Table S1:** List of oligonucleotides used in this work. **Table S2:** General Statistics of RNASeq data analysis. **Table S3:** List of differentially expressed genes (DEGs) in leaves of Low-Pi treated rice plants relative to control plants identified by RNASeq analysis. **Table S4:** Gene Ontology Enrichment analysis (GOEA) of genes that are regulated by treatment of rice plants with Low Pi in the category of Biological Processes. **Table S5:** Expression data of genes involved in Metabolism of Photosynthetic Pigments (Chlorophylls and carotenoids), and Chloroplast Biogenesis and Development in Low Pi-treated rice plants relative to Control plants. **Table S6:** Expression data of genes involved in leaf senescence in Low-Pi treated rice plants relative to Control plants. **Table S7:** Expression data of genes involved in Oxidative Stress in Low Pi-treated rice plants relative to Control plants. **Table S8:** Gene Ontology Enrichment analysis (GOEA) of genes that are regulated by treatment of rice plants with Low Pi in the category of Molecular Function. **Table S9:** Gene Ontology Enrichment analysis (GOEA) of genes that are regulated by treatment of rice plants with low Pi in the category of Cellular Component. **Table S10:** PANTHER enrichment analysis of Protein class in leaves of Low Pi plants relative to Control plants. **Table S11:** Expression data of genes involved in Phosphate homeostasis, signalling and uptake in Low Pi-treated rice plants relative to Control plants.

## References

Ames, B. N. (1966). Assay of Inorganic Phosphate, Total Phosphate and Phosphatases. Methods Enzymol. 8, 115–118. doi: 10.1016/0076-6879(66)08014-5

Barja, M. V., Ezquerro, M., Beretta, S., Diretto, G., Florez-Sarasa, I., Feixes, E., et al. (2021). Several geranylgeranyl diphosphate synthase isoforms supply metabolic substrates for carotenoid biosynthesis in tomato. New Phytologist 231, 255–272. doi: 10.1111/nph.17283

Bradford, M. M. (1976). A rapid and sensitive method for the quantitation of microgram quantities of protein utilizing the principle of protein-dye binding. Anal. Biochem. 72, 248–254. doi: 10.1016/0003-2697(76)90527-3

Bundó, M., Val-Torregrosa, B., Martín-Cardoso, H., Ribaya, M., Campos-Soriano, L., Bach-Pages, M., et al. (2024). Silencing Osa-miR827 via CRISPR/Cas9 protects rice against the blast fungus Magnaporthe oryzae. Plant Mol. Biol. 114, 105. doi: 10.1007/s11103-024-01496-z

Campo, S., Baldrich, P., Messeguer, J., Lalanne, E., Coca, M., and Segundo, B. S. (2014). Overexpression of a calcium-dependent protein kinase confers salt and drought tolerance in rice by preventing membrane lipid peroxidation. Plant Physiol. 165, 688–704. doi: 10.1104/pp.113.230268

Campos-Soriano, L., Bundó, M., Bach-Pages, M., Chiang, S.-F., Chiou, T.-J., and San Segundo, B. (2020). Phosphate excess increases susceptibility to pathogen infection in rice. Mol. Plant Pathol. 21, 555–570. doi: 10.1111/mpp.12916

Chen, S., Zhou, Y., Chen, Y., and Gu, J. (2018). fastp: an ultra-fast all-in-one FASTQ preprocessor. Bioinformatics 34, i884–i890. doi: 10.1093/bioinformatics/bty560

Chiang, C.-P., Yayen, J., and Chiou, T.-J. (2025). Mobilization and recycling of intracellular phosphorus in response to availability. Quantitative Plant Biology 6, e3. doi: 10.1017/qpb.2025.1

Chiou, T.-J., Aung, K., Lin, S.-I., Wu, C.-C., Chiang, S.-F., and Su, C. (2006). Regulation of Phosphate Homeostasis by MicroRNA in Arabidopsis. Plant Cell 18, 412–421. doi: 10.1105/tpc.105.038943

Falk, J., and Munné-Bosch, S. (2010). Tocochromanol functions in plants: antioxidation and beyond. J. Exp. Bot. 61, 1549–1566. doi: 10.1093/jxb/erq030

Gao, Q., Yang, Z., Zhou, Y., Yin, Z., Qiu, J., Liang, G., et al. (2012). Characterization of an Abc1 kinase family gene OsABC1-2 conferring enhanced tolerance to dark-induced stress in rice. Gene 498, 155–163. doi: 10.1016/j.gene.2012.02.017

George, T. S., Hinsinger, P., and Turner, B. L. (2016). Phosphorus in soils and plants – facing phosphorus scarcity. Plant Soil 401, 1–6. doi: 10.1007/s11104-016-2846-9

Gregersen, P. L., Holm, P. B., and Krupinska, K. (2008). Leaf senescence and nutrient remobilisation in barley and wheat. Plant Biol. 10, 37–49. doi: 10.1111/j.1438-8677.2008.00114.x

Guo, Y., Ren, G., Zhang, K., Li, Z., Miao, Y., and Guo, H. (2021). Leaf senescence: progression, regulation, and application. Molecular horticulture 1, 5. doi: 10.1186/s43897-021-00006-9

Hasanuzzaman, M., Bhuyan, M. H. M. B., Parvin, K., Bhuiyan, T. F., Anee, T. I., Nahar, K., et al. (2020). Regulation of ROS Metabolism in Plants under Environmental Stress: A Review of Recent Experimental Evidence. Int. J. Mol. Sci. 21, 8695. doi: 10.3390/ijms21228695

Jaisue, P., Daengngam, C., Pengphorm, P., Nutthapornnitchakul, S., Pinit, S., and Klinnawee, L. (2025). Enhanced Chlorophyll Accumulation is Early Response of Rice to Phosphorus Deficiency. Rice Sci. 32, 831–844. doi: 10.1016/j.rsci.2025.08.004

Lee, R., Wang, C., Huang, L., and Chen, S. G. (2001). Leaf senescence in rice plants: cloning and characterization of senescence up[regulated genes. J. Exp. Bot. 52, 1117–1121. doi: 10.1093/jexbot/52.358.1117

Lee, S., and Masclaux-Daubresse, C. (2021). Current Understanding of Leaf Senescence in Rice. Int. J. Mol. Sci. 22, 4515. doi: 10.3390/ijms22094515

Li, H., and Yu, C.-W. (2018). Chloroplast Galactolipids: The Link Between Photosynthesis, Chloroplast Shape, Jasmonates, Phosphate Starvation and Freezing Tolerance. Plant Cell Physiol. 59, 1128–1134. doi: 10.1093/pcp/pcy088

Liang, C., Wang, Y., Zhu, Y., Tang, J., Hu, B., Liu, L., et al. (2014). OsNAP connects abscisic acid and leaf senescence by fine-tuning abscisic acid biosynthesis and directly targeting senescence-associated genes in rice. Proc. Natl. Acad. Sci. U. S. A. 111, 10013–10018. doi: 10.1073/pnas.1321568111

Lichtenthaler, H. K., and Buschmann, C. (2001). Chlorophylls and Carotenoids: Measurement and Characterization by UV-VIS Spectroscopy. Current Protocols in Food Analytical Chemistry 1, F4.3.1–F4.3.8. doi: 10.1002/0471142913.faf0403s01

Lin, W.-Y., Huang, T.-K., and Chiou, T.-J. (2013). NITROGEN LIMITATION ADAPTATION, a Target of MicroRNA827, Mediates Degradation of Plasma Membrane–Localized Phosphate Transporters to Maintain Phosphate Homeostasis in Arabidopsis. Plant Cell 25, 4061–4074. doi: 10.1105/tpc.113.116012

Lu, Z., Ren, T., Li, Y., Cakmak, I., and Lu, J. (2025). Nutrient limitations on photosynthesis: from individual to combinational stresses. Trends Plant Sci. 30, 872–885. doi: 10.1016/j.tplants.2025.03.006

Martín-Cardoso, H., Bundó, M., Val-Torregrosa, B., and San Segundo, B. (2024). Phosphate accumulation in rice leaves promotes fungal pathogenicity and represses host immune responses during pathogen infection. Front. Plant Sci. 14, 1330349. doi: 10.3389/fpls.2023.1330349

Mayta, M. L., Hajirezaei, M.-R., Carrillo, N., and Lodeyro, A. F. (2019). Leaf Senescence: The Chloroplast Connection Comes of Age. Plants 8, 495. doi: 10.3390/plants8110495

Mi, H., Ebert, D., Muruganujan, A., Mills, C., Albou, L.-P., Mushayamaha, T., et al. (2021). PANTHER version 16: a revised family classification, tree-based classification tool, enhancer regions and extensive API. Nucleic Acids Res. 49, D394–D403. doi: 10.1093/nar/gkaa1106

Moustakas, M., Sperdouli, I., and Adamakis, I.-D. S. (2023). Editorial: Reactive oxygen species in chloroplasts and chloroplast antioxidants under abiotic stress. Front. Plant Sci. 14, 1208247. doi: 10.3389/fpls.2023.1208247

Nakano, Y., and Asada, K. (1981). Hydrogen Peroxide is Scavenged by Ascorbate-specific Peroxidase in Spinach Chloroplasts. Plant Cell Physiol. 22, 867–880. doi: 10.1093/oxfordjournals.pcp.a076232

Okazaki, Y., Otsuki, H., Narisawa, T., Kobayashi, M., Sawai, S., Kamide, Y., et al. (2013). A new class of plant lipid is essential for protection against phosphorus depletion. Nat. Commun. 4, 1510. doi: 10.1038/ncomms2512

Park, S.-Y., Yu, J.-W., Park, J.-S., Li, J., Yoo, S.-C., Lee, N.-Y., et al. (2007). The Senescence-Induced Staygreen Protein Regulates Chlorophyll Degradation. Plant Cell 19, 1649–1664. doi: 10.1105/tpc.106.044891

Poli, Y., Nallamothu, V., Balakrishnan, D., Ramesh, P., Desiraju, S., Mangrauthia, S. K., et al. (2018). Increased Catalase Activity and Maintenance of Photosystem II Distinguishes High-Yield Mutants From Low-Yield Mutants of Rice var. Nagina22 Under Low-Phosphorus Stress. Front. Plant Sci. 9, 1543. doi: 10.3389/fpls.2018.01543

Puga, M. I., Poza-Carrión, C., Martinez-Hevia, I., Perez-Liens, L., and Paz-Ares, J. (2024). Recent advances in research on phosphate starvation signaling in plants. J. Plant Res. 137, 315–330. doi: 10.1007/s10265-024-01545-0

Salvador-Guirao, R., Baldrich, P., Tomiyama, S., Hsing, Y.-I., Okada, K., and San Segundo, B. (2019). OsDCL1a activation impairs phytoalexin biosynthesis and compromises disease resistance in rice. Ann. Bot. 123, 79–93. doi: 10.1093/aob/mcy141

Scarpeci, T. E., Zanor, M. I., Carrillo, N., Mueller-Roeber, B., and Valle, E. M. (2008). Generation of superoxide anion in chloroplasts of Arabidopsis thaliana during active photosynthesis: a focus on rapidly induced genes. Plant Mol. Biol. 66, 361–378. doi: 10.1007/s11103-007-9274-4

Singh, S., Singh, A., and Nandi, A. K. (2016). The rice OsSAG12-2 gene codes for a functional protease that negatively regulates stress-induced cell death. J. Biosci. 41, 445–53. doi: 10.1007/s12038-016-9626-9

Sobieszczuk-Nowicka, E., Wrzesiński, T., Bagniewska-Zadworna, A., Kubala, S., Rucińska-Sobkowiak, R., Polcyn, W., et al. (2018). Physio-Genetic Dissection of Dark-Induced Leaf Senescence and Timing Its Reversal in Barley. Plant Physiol. 178, 654–671. doi: 10.1104/pp.18.00516

Supek, F., Bošnjak, M., Škunca, N., and Šmuc, T. (2011). REVIGO summarizes and visualizes long lists of gene ontology terms. PLoS One 6, e21800. doi: 10.1371/journal.pone.0021800

Tan, S., Sha, Y., Sun, L., and Li, Z. (2023). Abiotic Stress-Induced Leaf Senescence: Regulatory Mechanisms and Application. Int. J. Mol. Sci. 24, 11996. doi: 10.3390/ijms241511996

Versaw, W. K., and Harrison, M. J. (2002). A Chloroplast Phosphate Transporter, PHT2;1, Influences Allocation of Phosphate within the Plant and Phosphate-Starvation Responses. Plant Cell 14, 1751–1766. doi: 10.1105/tpc.002220

Wairich, A., Vitali, A., Adamski, J. M., Lopes, K. L., Duarte, G. L., Ponte, L. R., et al. (2023). Enhanced expression of OsNAC5 leads to up-regulation of OsNAC6 and changes rice (Oryza sativa L.) ionome. Genet. Mol. Biol. 46, e20220190. doi: 10.1590/1678-4685-gmb-2022-0190

Woo, H. R., Kim, H. J., Lim, P. O., and Nam, H. G. (2019). Leaf Senescence: Systems and Dynamics Aspects. Annu. Rev. Plant Biol. 70, 347–376. doi: 10.1146/annurev-arplant-050718-095859

Yang, S.-Y., Lin, W.-Y., Hsiao, Y.-M., and Chiou, T.-J. (2024). Milestones in understanding transport, sensing, and signaling of the plant nutrient phosphorus. Plant Cell 36, 1504–1523. doi: 10.1093/plcell/koad326

Zhang, J., Li, H., Xu, B., Li, J., and Huang, B. (2016). Exogenous Melatonin Suppresses Dark-Induced Leaf Senescence by Activating the Superoxide Dismutase-Catalase Antioxidant Pathway and Down-Regulating Chlorophyll Degradation in Excised Leaves of Perennial Ryegrass (Lolium perenne L.). Front. Plant Sci. 7, 1500. doi: 10.3389/fpls.2016.01500

Zhang, J.-F., Wang, Y.-Y., He, L., Yan, J.-Y., Liu, Y.-Y., Ruan, Z.-Y., et al. (2024a). PHR1 involved in the regulation of low phosphate-induced leaf senescence by modulating phosphorus homeostasis in Arabidopsis. Plant Cell Environ. 47, 799–816. doi: 10.1111/pce.14790

Zhang, Y., Wang, N., He, C., Gao, Z., and Chen, G. (2024b). Comparative transcriptome analysis reveals major genes, transcription factors and biosynthetic pathways associated with leaf senescence in rice under different nitrogen application. BMC Plant Biol. 24, 419. doi: 10.1186/s12870-024-05129-x

Zhao, Y., Zhang, Y., Li, S., Tan, S., Cao, J., Wang, H.-L., et al. (2024). Leaf Senescence Database v5.0: A Comprehensive Repository for Facilitating Plant Senescence Research. J. Mol. Biol. 436, 168530. doi: 10.1016/j.jmb.2024.168530

