## Supplementary Figures for "Molecular basis of delayed leaf senescence induced by short-term treatment with low phosphate in rice"

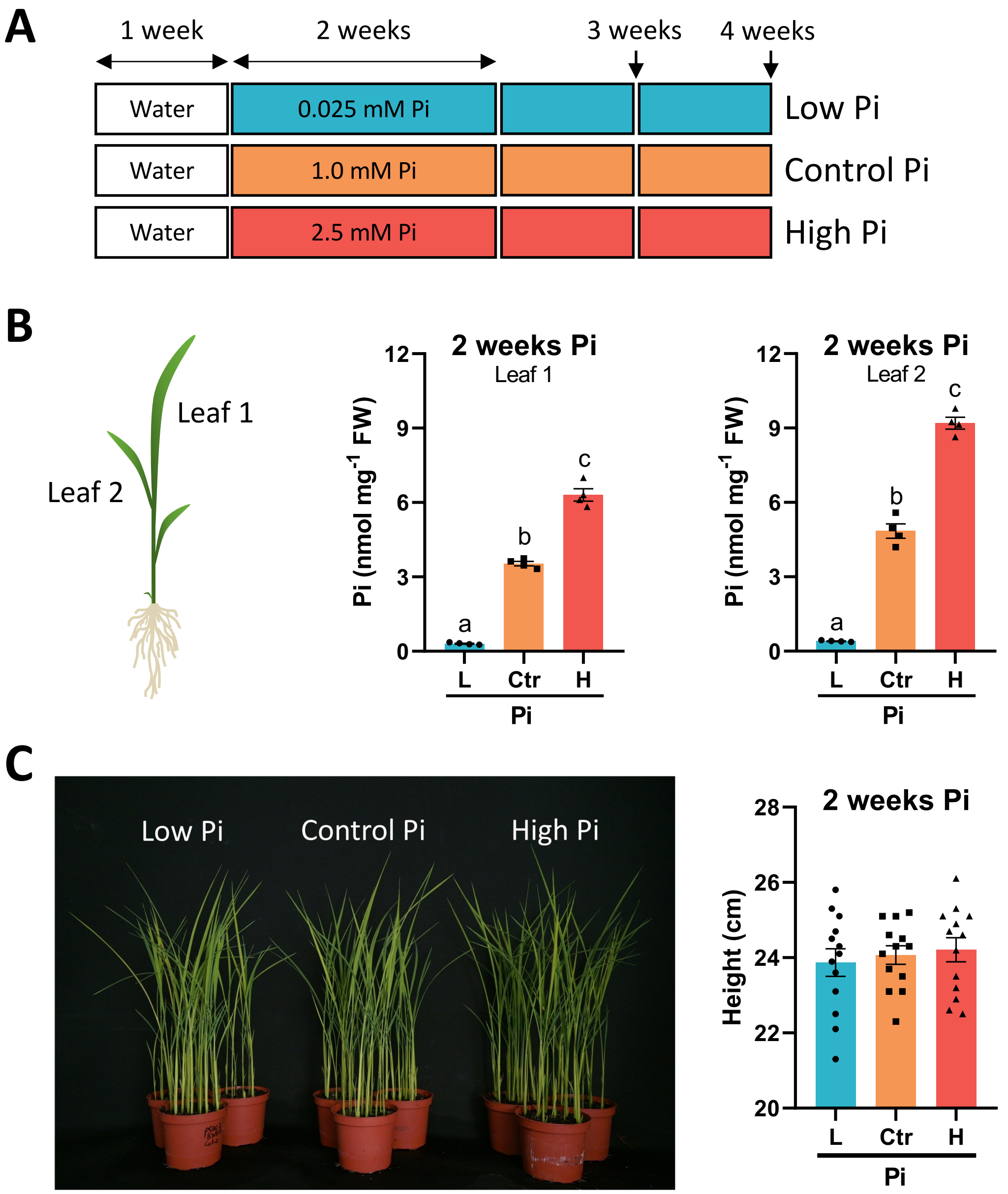


**Figure S1**. **Conditions used for Pi treatment of rice seedlings**. (**A**) Seedlings were pregerminated in water for 7 days, transplanted to 50% turface and vermiculite combined with 50% quartz sand, and then fertilized for the desired period of time with nutrient solution containing 0.025 mM, 1 mM or 2.5 mM Pi for 2 weeks (Low Pi, Control, and High Pi plants, respectively). (**B**) Leaves were numbered starting from the youngest, fully expanded leaf (Leaf 1). Middle and right panels, Pi content in leaves (Leaf 1, Leaf 2) at 2 weeks of Pi treatment. Bars represent mean ± SEM of 4 biological replicates (each replicate from a pool of 4 plants). (**C**) Appearance and height of rice seedlings that have been treated with Pi for 14 days (N = 15).

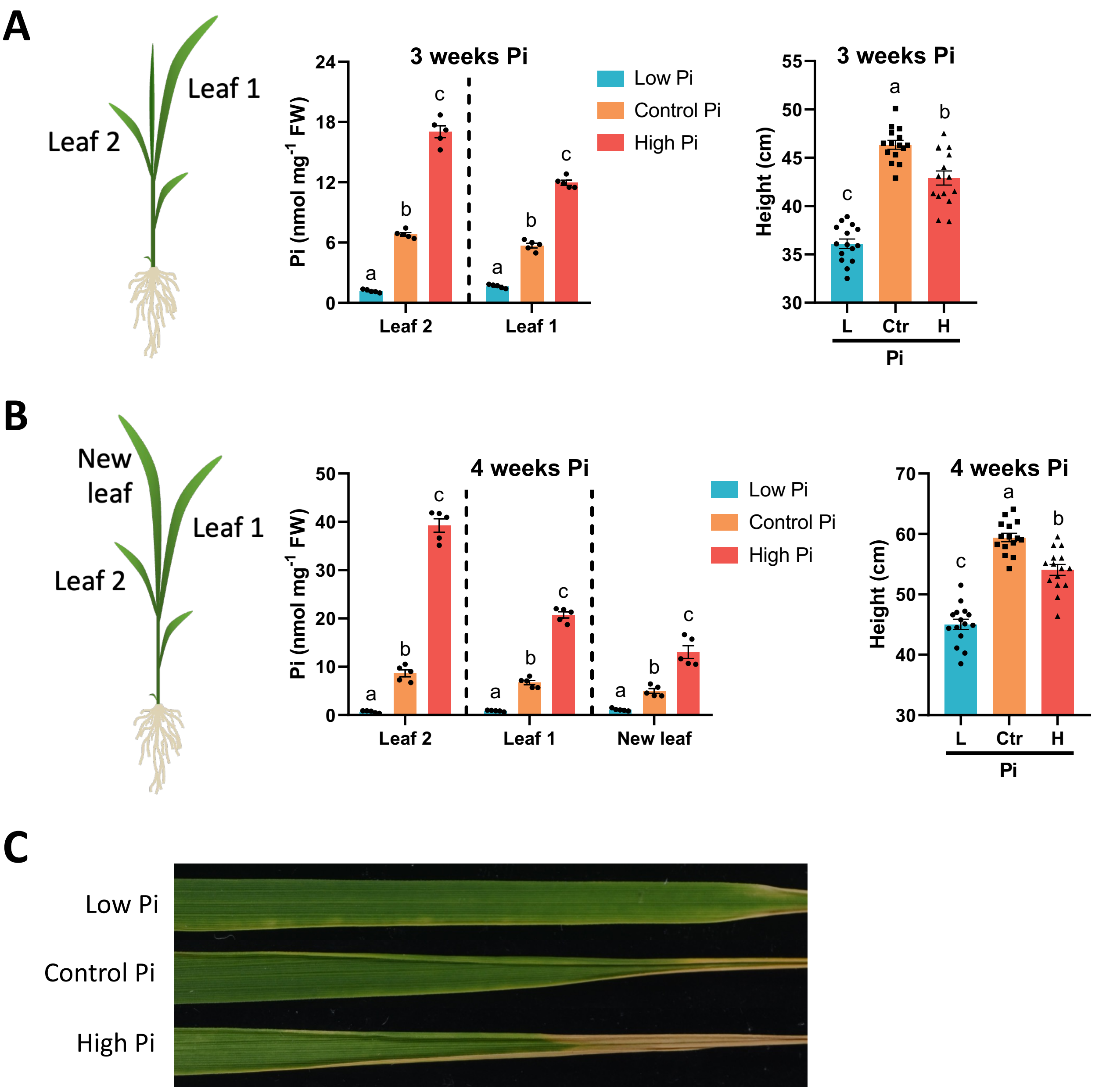


**Figure S2. Effect of treatment of rice seedlings with Pi for 3 and 4 weeks**. (**A** and **B**) Pi content (N = 5) and height (N = 15) in leaves at 3 weeks (**A**) or 4 weeks (**B**) of Pi treatment. (**C**) Leaf tip necrosis at 4 weeks of Pi treatment.

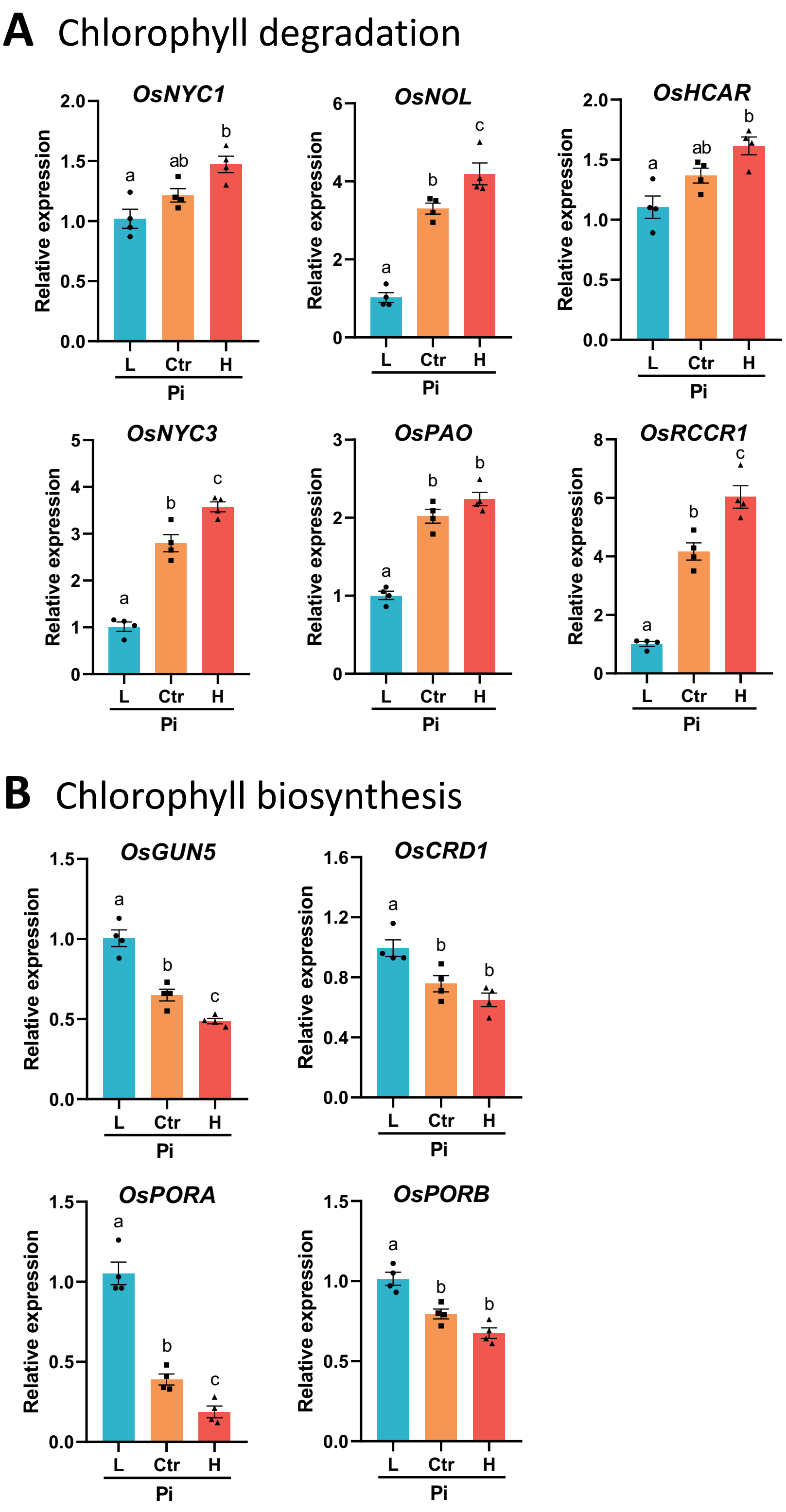


**Figure S3. Expression of genes involved in chlorophyll metabolism in leaves of rice plants that have been grown under different Pi conditions**. Expression was determined by RT-qPCR using the rice *Ubiquitin 1* (*OsUbi1*) gene to normalize transcript levels. (**A**) Genes involved in chlorophyl degradation. (**B**) Genes involved in chlorophyll biosynthesis. Bars represent mean ± SEM of 4 biological replicates, each one from a pool of 4 different plants. Statistically significant differences were determined by one-way ANOVA (different letters indicate significant differences among Pi conditions in each leaf).

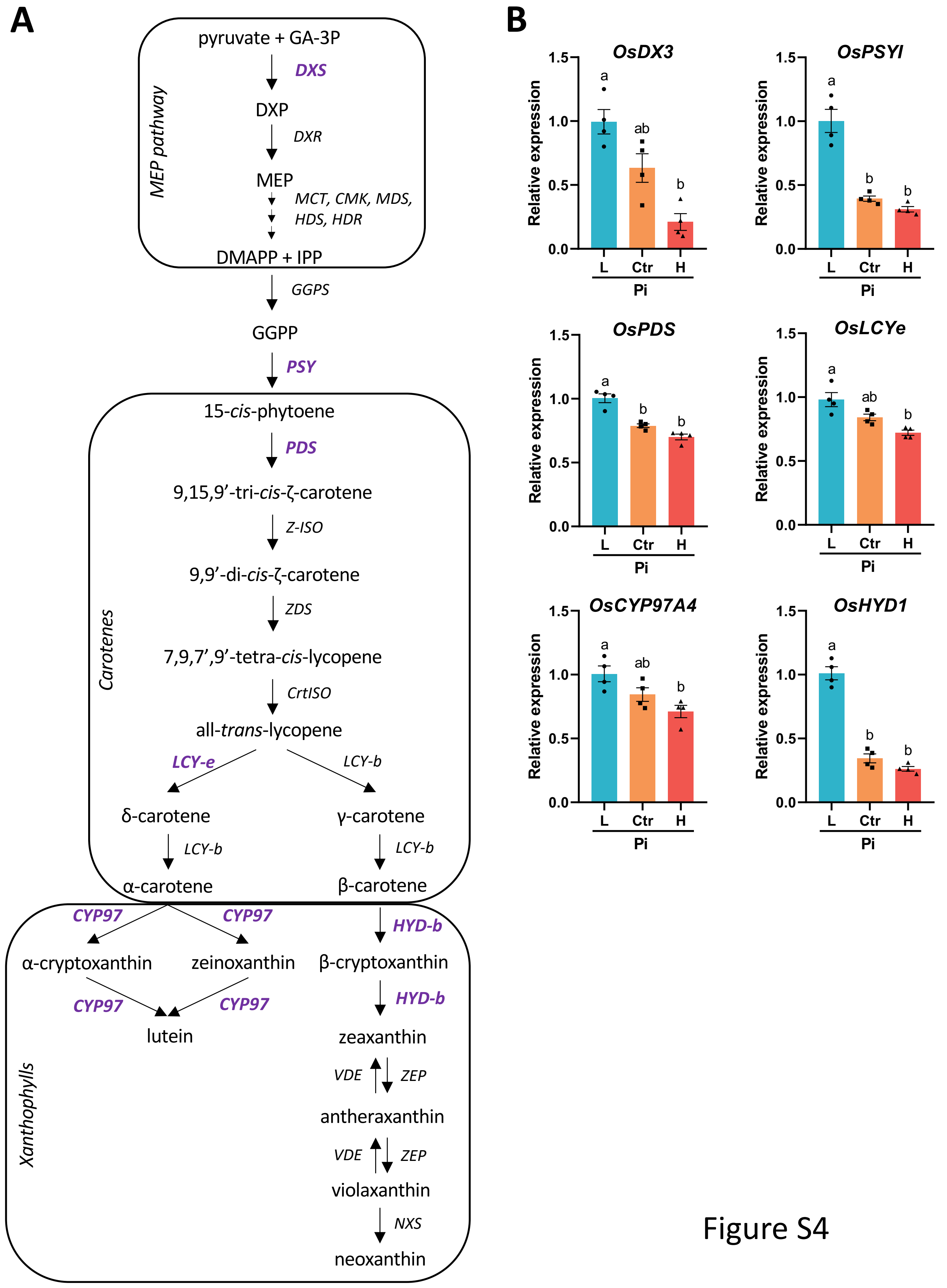

A
pyruvate + GA-3P
DXS
DXP
DXR
MEP
MCT, CMK, MDS,
HDS, HDR
DMAPP + IPP
MEP pathway
GGPS
GGPP
PSY
15-cis-phytoene
PDS
9,15,9'-tri-cis-ζ-carotene
Z-ISO
9,9'-di-cis-ζ-carotene
ZDS
7,9,7',9'-tetra-cis-lycopene
CrtISO
all-trans-lycopene
LCY-e
LCY-b
δ-carotene
γ-carotene
LCY-b
LCY-b
α-carotene
β-carotene
Carotenes
CYP97
CYP97
HYD-b
α-cryptoxanthin
zeinoxanthin
β-cryptoxanthin
CYP97
CYP97
HYD-b
lutein
zeaxanthin
VDE
ZEP
antheraxanthin
VDE
ZEP
violaxanthin
NXS
neoxanthin
Xanthophylls
B
OsDX3
OsPSYI
OsPDS
OsLCYe
OsCYP97A4
OsHYD1
Relative expression
1.5
1.0
0.5
0.0
L
Ctr
H
Pi
a
ab
b


Figure S4

**Figure S4**. **Expression of genes involved in carotenoid biosynthesis in rice plants grown under different Pi supply**. (**A**) Schematic view of the MEP (methylerythritol phosphate) pathway that produces the building blocks (DMAPP and IPP) used in carotenoid biosynthesis. Genes whose expression was examined in this study are shown in bold. . (**B**) Expression of genes involved in carotenoid biosynthesis determined by RT-qPCR. Bars represent mean ± SEM of 4 biological replicates, each one from a pool of 4 different plants. Statistically significant differences were determined by one-way ANOVA (different letters indicate significant differences among Pi conditions in each leaf). CMK, 4-(cytidine 5'-diphospho)-2-C-methyl-D-erythritol kinase; CrtISO, carotene *cis-trans*-isomerase; CYP97, cytochrome P450 carotene hydroxylases; DMAPP, dimethylallyl diphosphate; DXP, deoxyxylulose 5-phosphate; DXR, deoxyxylulose 5-phosphate reductoisomerase; GA-3P, d-fluceraldehyde-3-phosphate; GGPP, geranylgeranyl-diphosphate; GGPS, geranylgeranyl diphosphate synthase; HDR, 1-hydroxy-2-methyl-2-(E)-butenyl 4-diphosphate reductase; HDS, 1-hydroxy-2-methyl-butenyl 4-diphosphate synthase; HYD-b, carotene β-ring hydroxylase 1; IPP, isopentenyl diphosphate; LCY, lycopene cyclase; MCT, 4-(cytidine 5'-diphospho)-2-C-methyl-D-erythritol synthase; MDS, 2-C-methyl-D-erythritol 2,4-cyclodiphosphate synthase; MEP, methylerythritol 4-phosphate; NXS, neoxanthin synthase; PDS, phytoene desaturase; PSY, phytoene synthase; VDE, violaxanthin de-epoxidase; Z-ISO, ζ-carotene isomerase; ZDS, ζ-carotene desaturase; ZEP, zeaxanthin epoxidase.

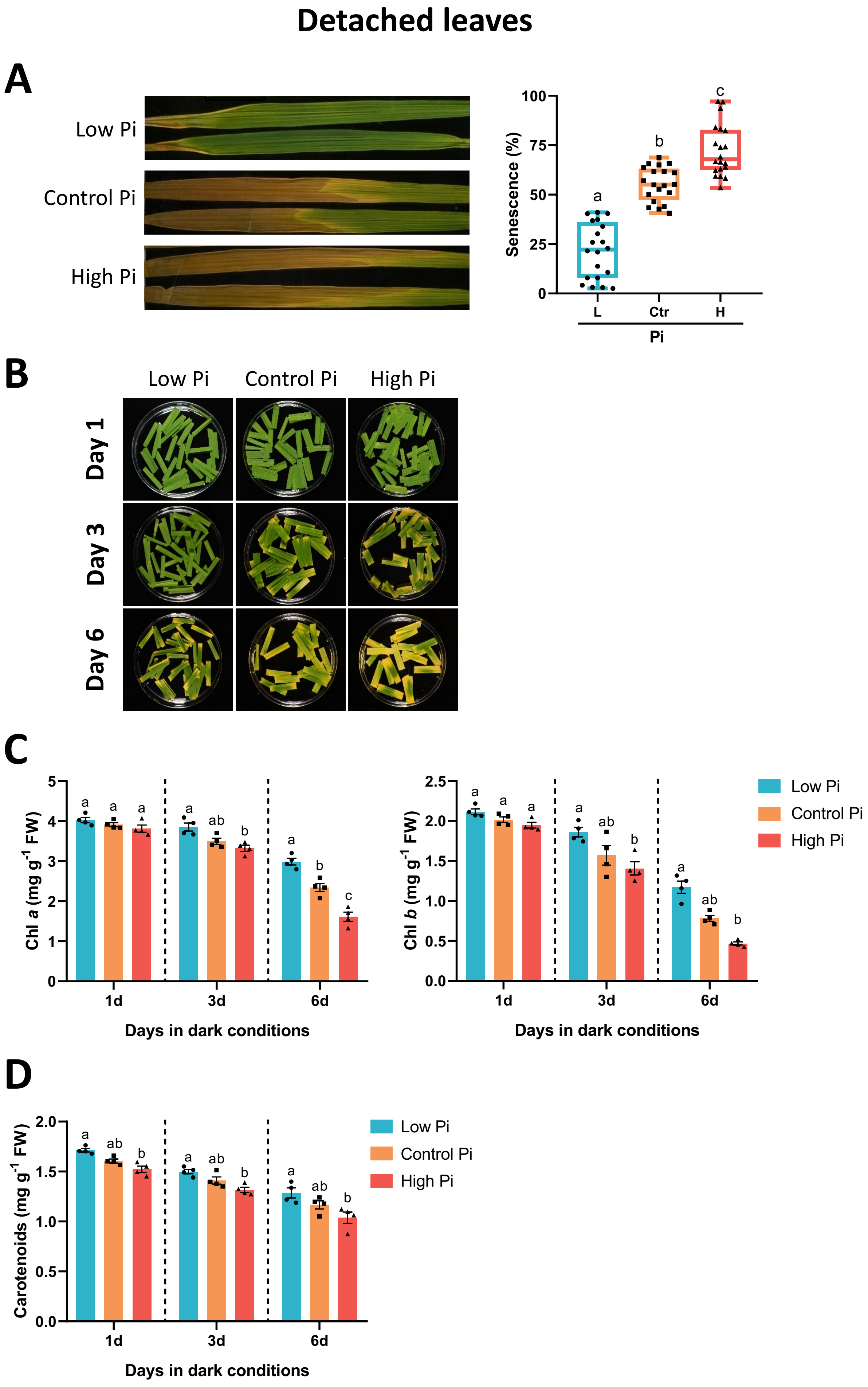
Detached leaves
A
Low Pi
Control Pi
High Pi
Senescence (%)
100
75
50
25
0
a
b
c
L
Ctr
H
Pi
B
Low Pi
Control Pi
High Pi
Day 1
Day 3
Day 6
C
Chl a (mg g-1 FW)
Chl b (mg g-1 FW)
1d
3d
6d
Days in dark conditions
Low Pi
Control Pi
High Pi
D
Carotenoids (mg g-1 FW)
1d
3d
6d
Days in dark conditions
Low Pi
Control Pi
High Pi

Figure S5

**Figure S5**. **Senescence phenotype of detached leaves from rice seedlings grown under increasing Pi supply for 2 weeks**. Leaves, either entire leaves or leaf segments, were then subjected to dark treatment. Images from Leaf 1 are shown (similar results were obtained with leaf sections from Leaf 2. (**A**) Appearance of entire leaves from Pi-treated seedlings at 9 days of dark treatment. Right panel, percentage of senescence determined by image analysis using the software APS Assess 2.0. The senescence level is represented as the % of the leaf area showing senescence relative to the total leaf area. Box plots represent median and data distribution (N = 20 leaves, each condition). (**B**) Appearance of leaf sections at 1 day, 3 days or 6 days of dark treatment. Each condition consisted of sections from leaves of 6 independent plants. (**C**) Chlorophyll content analysis, Chl *a* and Chl *b*. (**D**) Content of total carotenoids. Bars in **C** and **D** represent mean ± SEM of 4 biological replicates, each one from a pool of 4 different plants. Results are expressed as mg/g of fresh weight. Statistically significant differences were determined by one-way ANOVA (different letters indicate significant differences among Pi conditions). Three independent experiments were conducted with similar results.

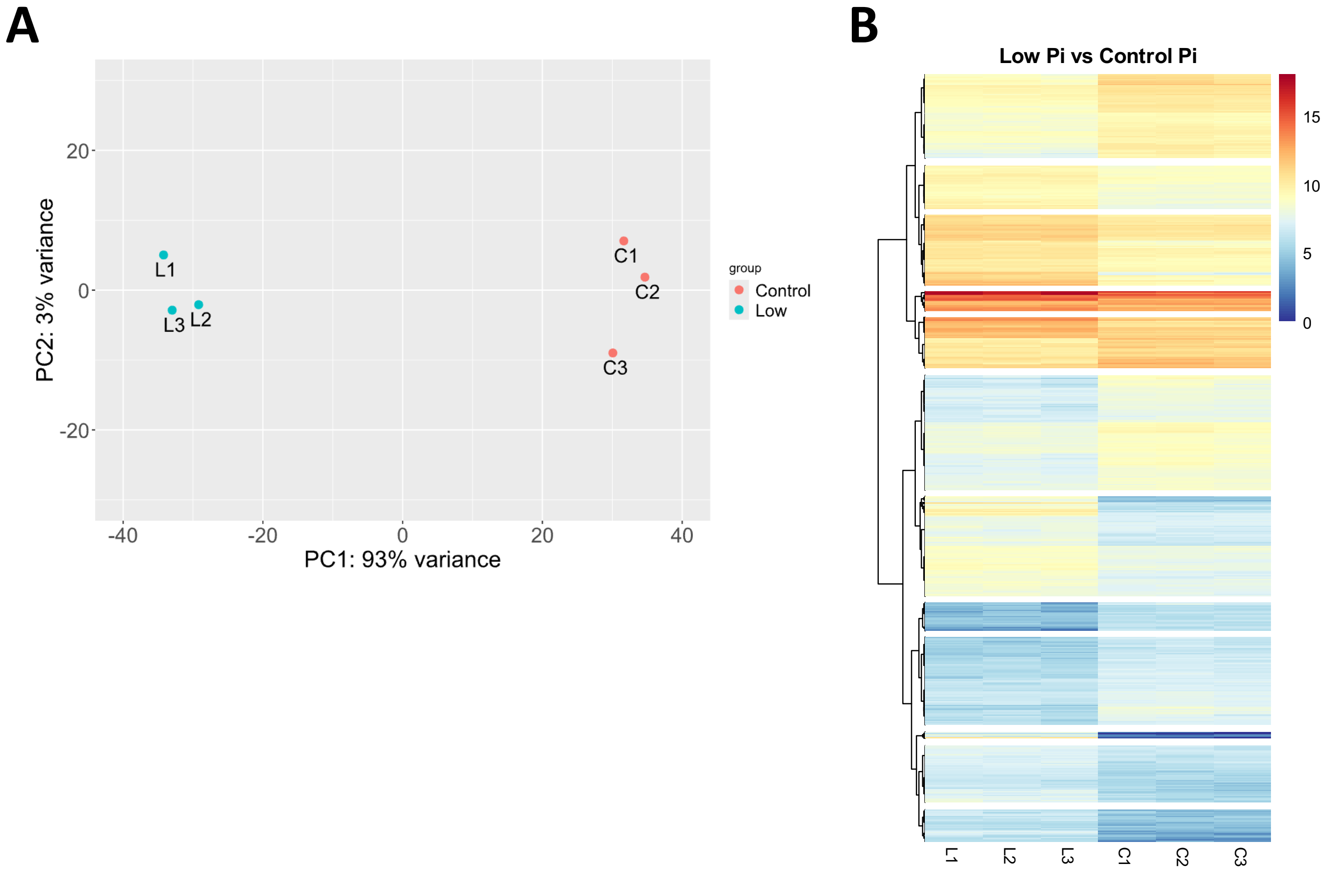


**Figure S6**. **Analysis of RNASeq data from leaves of Low-Pi and Control plants**. **(A)** Principal component analysis (PCA) analysis of RNA-Seq data to visualize sample to sample variation. The PCA plot depicted clear clustering of samples from control plants (red dots) and samples from plants treated with low Pi (blue dots). **(B)** Hierarchical cluster of gene expression in leaves of Low-Pi plants based on the $Log_2$ of normalized counts. The colour scale ranges from blue (low expression) to red (high expression).

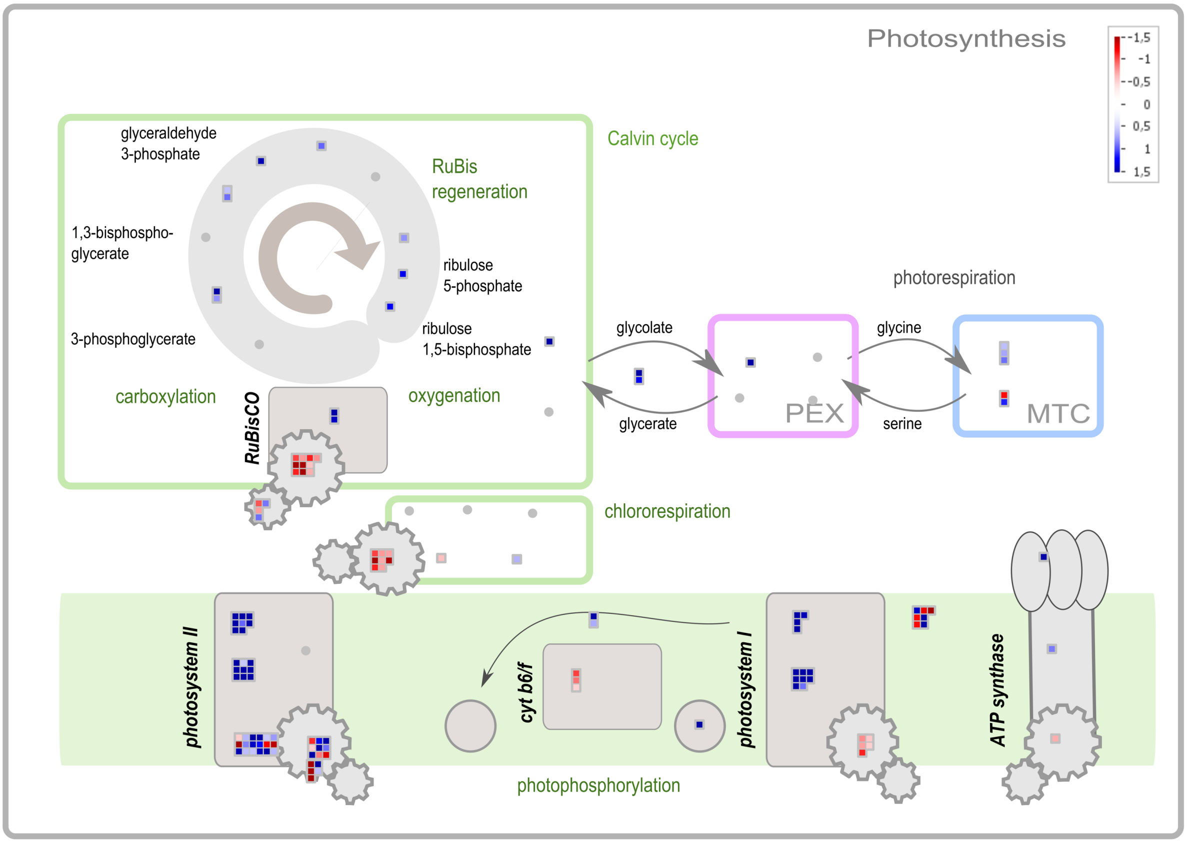


**Figure S7**. **MapMan overview of Photosynthesis**. The colour key represents the log2FC (values scaled to 1.5 to -1.5). Red represents the down-regulation and blue represents the upregulation of rice genes in leaves of Low-Pi rice plants relative to control plants. PEX, peroxisome; MTC, mitochondrion.

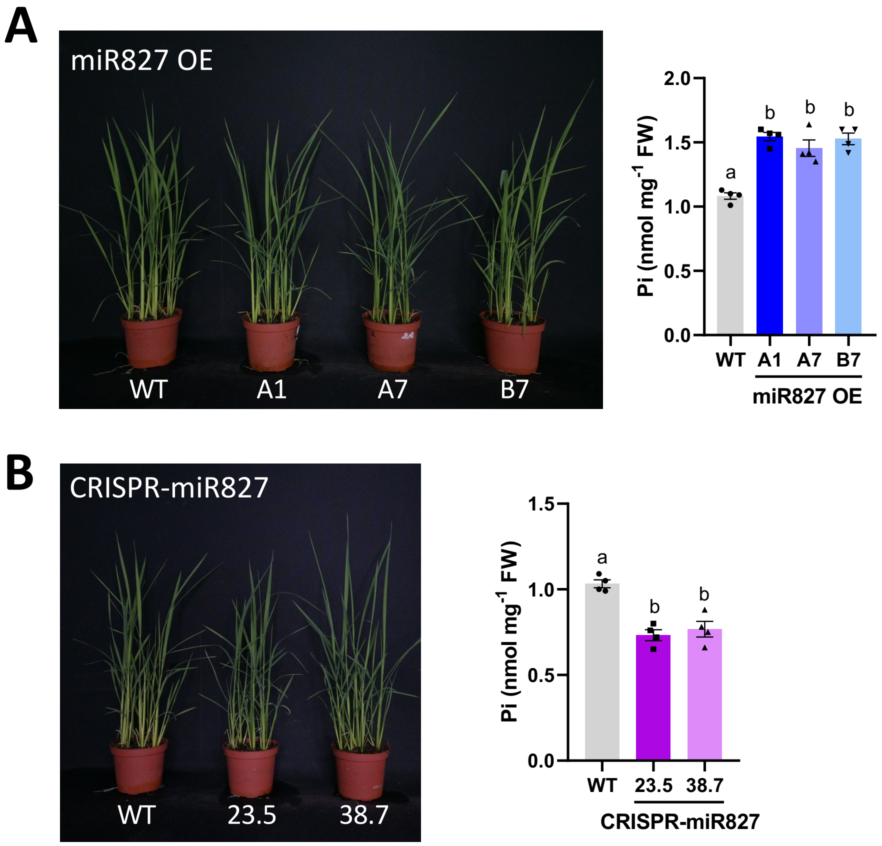


**Figure S8**. **Phenotype of miR827 overexpressor and CRISPR/Cas9-edited miR827 rice plants**. Three independent experiments were conducted with similar results. Statistically significant differences were determined by one-way ANOVA (different letters indicate significant differences among Pi conditions). **(A** and **B)** Appearance of three-week-old miR827OE plants (lines A1, A7 and B7) **(A)** and CRISPR-miR827 plants (lines 23.5 and 38.7) **(B)**. Right panels show the Pi content (free Pi) of leaves (Leaf 1) from miR827 OE and CRISPR-miR827 plants. Bars represent mean ± SEM of 4 biological replicates (each replicate from a pool of 4 plants).

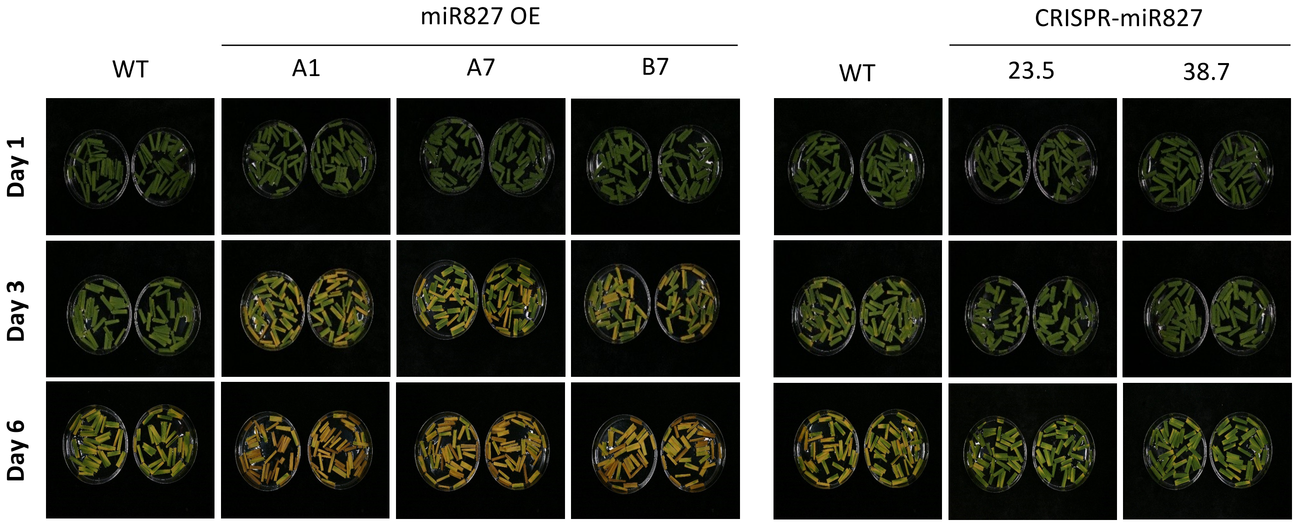


**Figure S9**. **Senescence phenotype of detached leaf sections from miR827 OE or CRISPR-miR827 plants subjected to darkness during 1, 3 and 6 days**. Representative images of leaves from seedlings at the 3-4 leaf stage are shown. Each condition consisted of sections from leaves of 6 independent plants per genotype.

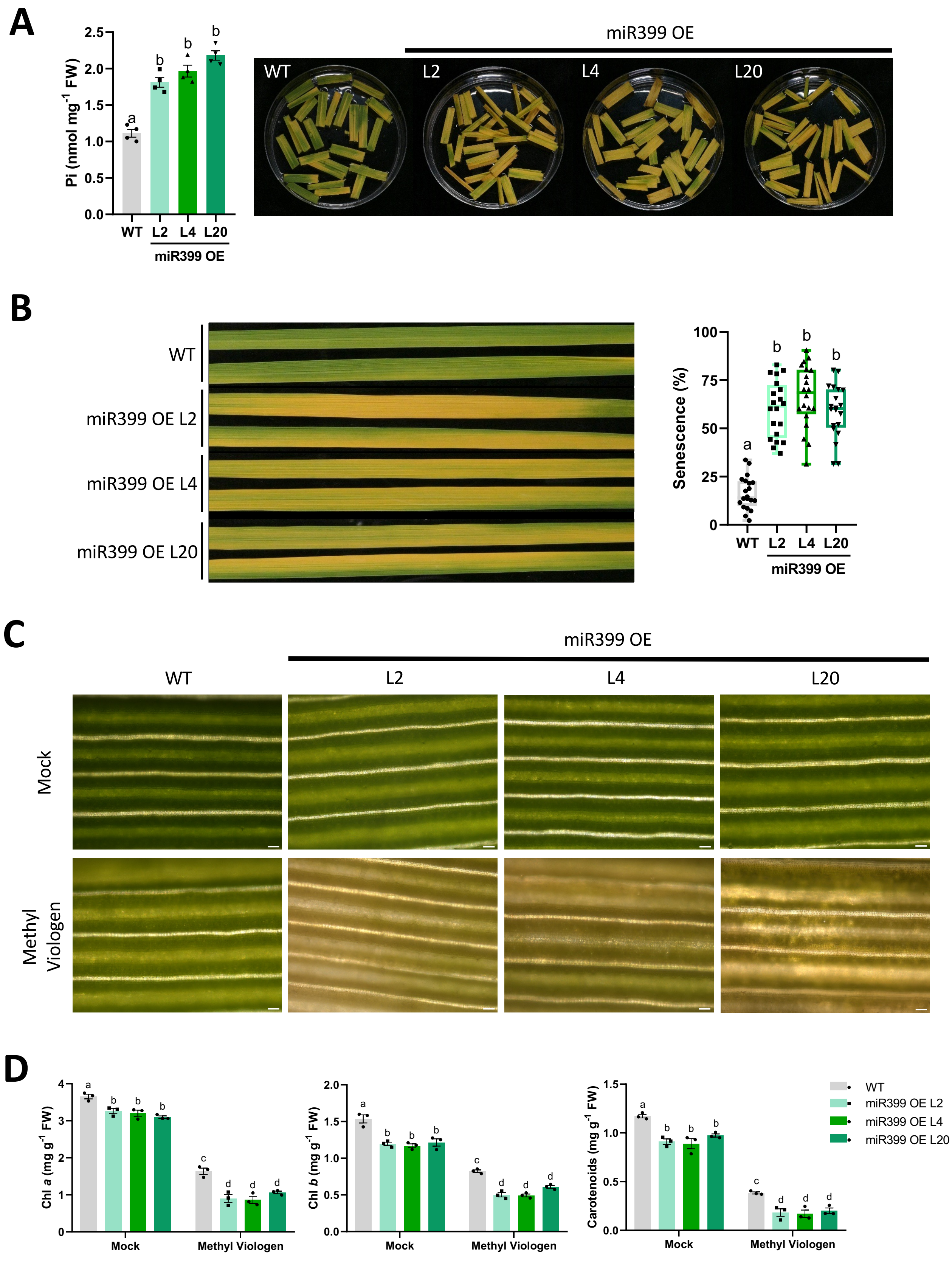
A
Pi (nmol mg-1 FW)
2.5
2.0
1.5
1.0
0.5
0.0
a
b
b
b
WT
L2
L4
L20
miR399 OE
miR399 OE
WT
L2
L4
L20
B
WT
miR399 OE L2
miR399 OE L4
miR399 OE L20
Senescence (%)
100
75
50
25
0
a
b
b
b
WT
L2
L4
L20
miR399 OE
C
miR399 OE
WT
L2
L4
L20
Mock
Methyl Viologen
D
Chl a (mg g-1 FW)
4
3
2
1
0
a
b
b
b
c
d
d
d
Mock
Methyl Viologen
Chl b (mg g-1 FW)
2.0
1.5
1.0
0.5
0.0
a
b
b
b
c
d
d
d
Mock
Methyl Viologen
Carotenoids (mg g-1 FW)
1.5
1.0
0.5
0.0
a
b
b
b
c
d
d
d
Mock
Methyl Viologen
WT
miR399 OE L2
miR399 OE L4
miR399 OE L20

Figure S10

**Figure S10**. **Senescence phenotype of leaves from rice plants overexpressing miR399 (lines L2, L4 and L20)**. Two independent experiments were conducted with similar results. Statistically significant differences were determined by one-way ANOVA (different letters indicate significant differences among genotypes). **(A)** Pi content in leaves (Leaf 1) of wild-type (WT) and miR399 OE plants (left panel). Senescence phenotype of leaves (Leaf 1) during DILS, after 6 days of darkness (right panel). **(B)** Senescence phenotype of entire leaves (Leaf 1) of miR399 OE plants during DILS (9 days of dark treatment). Quantification of senescence regions (N = 20) by image analysis using the software APS Assess 2.0 (right panel). **(C)** Effect of treatment with methyl viologen (MV) in WT and miR399 overexpressor plants. Scale bar = 50 µm. **(D)** Content of chlorophylls (Chl *a*, Chl *b*) and carotenoids in MV-treated rice leaves of WT and miR399 OE plants. Bars represent mean ± SEM of 3 biological replicates, each one from a pool of 4 different plants.

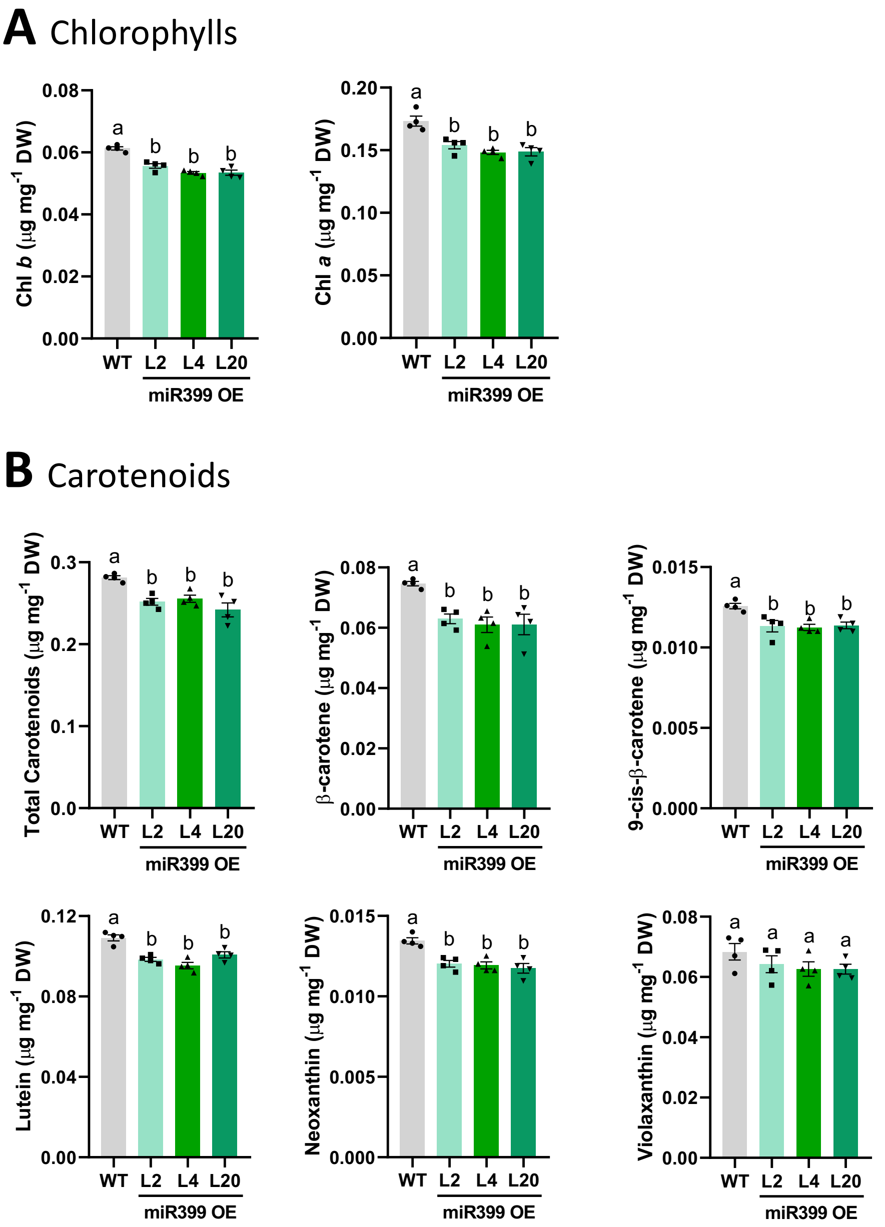


**Figure S11**. **Content of photosynthetic pigments, chlorophylls and carotenoids, in leaves of miR399 OE rice plants**. Leaves from plants at the 3-4 leaf stage (Leaf 1, youngest leaf) were used for HPLC analysis of photosynthetic pigments. Bars represent mean ± SEM of 4 biological replicates, each one from a pool of 4 different plants. Three independent experiments were conducted. Statistically significant differences were determined by one-way ANOVA (different letters indicate significant differences among Pi conditions). **(A)** Chlorophyll *a* and Chlorophyll *b* content in miR399 OE plants. **(B)** Carotenoid content (total carotenoids, lutein, 9-cis-β-carotene, β-carotene, violaxanthin and neoxanthin) in miR399 overexpressor rice plants.
